# Genetic and Pharmacologic Targeting of Eya3 in Macrophages Drives Anti-Tumor Immunity in Triple-Negative Breast Cancer

**DOI:** 10.64898/2026.07.22.739910

**Authors:** Kaiah M. Fields, Gabriela Terue Rizzo Kodama, Etienne Danis, Jessica G. Olivas-Corral, Sheera R. Rosenbaum, Uma Kantheti, Goksu Sarioglu, Erin Citarella, Lars Wick, Kate Matlin, Brooke Aloe, Emmy G. Hawkins, Arthur R. Wolin, Alexander LaVeck, Connor J. Hughes, Xiang Wang, Rui Zhao, Beth A. J. Tamburini, Gina Bouchard, Jill E. Slansky, Heide L. Ford

**Affiliations:** Department of Pharmacology, University of Colorado Anschutz, Aurora, CO, USA; Molecular Biology Program, University of Colorado Anschutz, Aurora, CO, USA; Department of Pharmaceutical Sciences, University of Colorado Anschutz, Aurora, CO, USA; Toxicology Graduate Program, University of Colorado Anschutz, Aurora, CO, USA; Department of Biomedical Informatics, University of Colorado Anschutz, Aurora, CO, USA; University of Colorado Cancer Center University of Colorado Anschutz, Aurora, CO, USA; Immunology Graduate Program, University of Colorado Anschutz, Aurora, CO, United States; Division of Gastroenterology and Hepatology, Department of Medicine, University of Colorado Anschutz, Aurora, CO, United States; Medical Scientist Training Program, University of Colorado Anschutz, Aurora, CO, USA; Cancer Biology Program, University of Colorado Anschutz, Aurora, CO, United States; Department of Biochemistry and Molecular Genetics, University of Colorado Anschutz, Aurora, CO, USA; Department of Chemistry, University of Colorado Boulder, Boulder, CO, USA; Department of Immunology and Microbiology, University of Colorado Anschutz, Aurora, CO, United States

## Abstract

Triple-negative breast cancer (TNBC) is an aggressive form of breast cancer that remains difficult to treat despite its relatively high immunogenicity, as tumors frequently evade immune destruction through mechanisms that are not yet fully understood. Here, we discover a previously unrecognized role for Eya3 within macrophages in the tumor immune microenvironment, where its expression is elevated and correlated with increased aggressiveness in TNBC. Using macrophage-targeted Eya3 knockdown and conditional knockout mouse models, we show that Eya3 depletion induces coordinated transcriptional and functional changes in macrophages, enhancing migration, antigen processing, and inflammatory signaling associated with anti-tumor immunity. Strikingly, macrophage-targeted deletion of Eya3 reprograms the immune response in TNBC, increasing CD8+ T cell infiltration, suppressing primary tumor growth, and prolonging survival. Pharmacologic inhibition of Eya1-3 tyrosine phosphatase activity with a newly derived allosteric inhibitor, LG1-34, mirrors this effect, dramatically reducing primary TNBC growth through immune-mediated mechanisms that likely act both though targeting tumor and immune cells. Importantly, TNBC patients treated with pembrolizumab exhibit high expression of Eya3 in both tumor and immune compartments, and increased macrophage specific Eya3 expression correlates with worsened response to anti-PD1 therapy. These findings identify Eya3 as a macrophage-intrinsic checkpoint on anti-tumor immunity, identifying a new potential vulnerability in TNBC.

**Significance Statement:** Targeting Eya3 Tyr phosphatase activity in tumor-associated macrophages reprograms the TNBC immune microenvironment and restores anti-tumor immunity, identifying a new potential therapeutic vulnerability in a cancer that has limited treatment targeted options.

## Introduction

Breast cancer (BC) is the most commonly diagnosed cancer among women, and the second leading cause of cancer death, with 15-20% of cases classified as triple negative breast cancer (TNBC)[1]. TNBC is an aggressive, heterogeneous, and highly metastatic subtype that has fewer treatment options than other invasive cancers due to the absence/low levels of targetable molecules such as estrogen and progesterone receptors, as well as HER2[2]. Current treatment options for TNBC include surgical resection, chemotherapy, and radiation[3]. PARP inhibitors and immune checkpoint inhibitors are approved in combination with chemotherapy for a subset of TNBC patients, but targeted treatment options remain limited and incompletely effective for most patients [3, 4].

TNBC is further characterized by its genomic instability and mutational load, leading to increased neoantigen production that the immune system can react to as “non-self”[5, 6]. Accordingly, TNBC is characterized as the most immunogenic of the breast tumor subtypes, yet TNBC cells often avoid immune destruction, thus counteracting the cytotoxic efforts of the immune system[7]. Work towards improving immunotherapy in TNBC is therefore largely focused on finding additional targets on cancer cells or immune cells that when targeted, improve the immune system’s ability to recognize and eliminate cancer cells[4, 8–10].

In breast and other cancers, embryonic signaling programs are often activated outside of their temporally appropriate context, contributing to tumor progression by reactivating pathways important for organ development[11–14]. While such oncofetal signaling is primarily thought to promote tumor development through tumor cell intrinsic mechanisms, embryonic programs can also aid growth by suppressing the tumor immune microenvironment (TIME), similar to what is observed in the crosstalk between maternal and embryonic cells that results in maternal-fetal immune tolerance[15–18]. We previously found that Eya3, a member of the multifunctional Eyes absent family (Eya1-4), is an example of a developmental protein that is upregulated in TNBC and promotes primary and metastatic tumor growth in part by suppressing adaptive and innate cytotoxic immune responses[19, 20].

Eyes absent (Eya) was first discovered in *Drosophila* as necessary for eye development (loss of the *Drosophila eya* gene results in no eyes, thus the designation of “eyes absent”)[21–23]. Eya proteins are critical for the development of multiple organs in mammals, including the eye, ear, muscle, kidney, and heart, amongst others [22, 24–27]. They have three discrete biochemical activities-intrinsic tyrosine (Tyr) phosphatase activity, associated serine/threonine (Ser/Thr) phosphatase activity via interaction with protein phosphatase 2A (PP2A), and transactivation activity when bound to the Six family of transcription factors (TFs)[28]. The multifunctional, pro-growth nature of Eya proteins have implicated them in numerous cancers, as their upregulation leads to tumor cell autonomous effects including increased proliferation, migration and invasion [28–32], as well as non-tumor cell autonomous effects, particularly on the TIME[19, 20]. For example, suppression of CD8+ T cells in TNBC primary tumors has been attributed to upregulation of tumor cell PD-L1 via c-Myc stabilization, an effect that is mediated by Eya3’s associated Ser/Thr phosphatase activity[19, 33], and suppression of cytotoxic NK cell activity at the lung premetastatic niche in TNBC is attributed to Eya3’s regulation of NFkB signaling[20].

CD8+ T cells and NK cells are two of several immune cell types that collectively contribute to the tumor immune response. The immune system is comprised of innate (macrophages, dendritic cells, NK cells, etc.) and adaptive immune cells (T cells, B cells), with innate immune cells acting as the first responders during immune challenges, including cancer. Cancer cells and immune cells have a reciprocal relationship in which they respond to and influence each other’s phenotypes[34]. While cancer cell plasticity has been one of the greatest challenges to targeted therapy, the mutual plasticity between cancer and immune cells offers a therapeutic niche that can be leveraged to restore immune elimination of tumors[34]. Macrophages, an innate immune cell population, are an intriguing therapeutic target in TNBC, as they can make up to 50% of the tumor volume, and while often existing in a “pro-tumor” state during established disease, they are extraordinarily responsive to their environment and can shift from having pro-tumor to anti-tumor functions depending on the cues they receive[8, 35–45]. Given the large effect of tumor-expressed Eya3 on the TIME, and the fact that Eya3 regulates critical signaling pathways in tumor cells that are also important in immune cells (Myc, NFkB)[19, 20], we asked whether Eya proteins may play a role in the immune cells themselves, particularly during tumor progression.

In this study, we show that the developmental protein, Eya3, is critical not only in TNBC cells, but also for macrophage function, altering the TIME. Analysis of mouse and human gene expression datasets reveals that Eya3 is expressed across immune populations, with increased expression in tumor-associated macrophages. Eya3 loss, via knockdown in macrophage cell lines (RAW264.7 and J774A.1) or knockout in primary bone marrow derived macrophages (BMDMs) from a macrophage-targeted conditional knockout mouse (Eya3fl/fl;MafB-Cre) we developed, enhances macrophage migration and antigen processing, with RNA-seq confirming transcriptional reprogramming towards anti-tumor macrophage signatures. *In vivo*, macrophage-targeted Eya3 deletion reduces primary TNBC growth, improves survival, and enhances CD8+ T cell infiltration. Pharmacologic inhibition using a newly developed small molecule Eya1-3 tyrosine phosphatase inhibitor, LG1-34[46], phenocopies these effects, boosting macrophage migration and antigen processing, while also reducing PD-L1 surface expression on E0771 tumor cells, consistent with prior work on tumor-intrinsic Eya3 knockdown (KD)[19]. In C57BL/6 E0771 tumor-bearing mice, LG1-34 treatment recapitulates the effect observed with Eya3 KO, enhancing CD8+ T cell infiltration, significantly reducing tumor growth, and extending survival. These effects are lost in immune compromised NOD scid gamma (NSG) mice, confirming that efficacy of Eya3 targeting requires an intact immune system. Further, we show that high macrophage Eya3 expression correlates with worse response to immunotherapy, and high Eya1 and 3 together correlate with decreased probability of survival after immunotherapy. Together, these findings underscore a previously unrecognized role for Eya3 in macrophages, and provide proof-of-concept that targeting Eya3 in TNBC may be a powerful way to reverse immune suppression and promote immune-mediated tumor elimination.

## Results

### Eya3 is expressed in macrophages, enriched in tumor-associated macrophages, and its macrophage specific expression correlates with increased TNBC grade

We previously reported that tumor expressed Eya3 promotes primary tumor growth and metastasis in TNBC by suppressing CD8+ T cell- and NK cell-mediated anti-tumor immune responses [20, 47]. In the TNBC context, we found that Eya3 regulates these immune responses by enhancing two critical signaling pathways that are prevalent in both tumor cells and immune cells within the TIME: Myc and NFkB[19, 20]. Because these pathways play an important role in immune cells, we examined whether Eya3 may also function in the TIME. To this end, we first examined whether Eya3 is expressed in immune cells using publicly available datasets. RNA-seq data from the Immunological Genome Project (Immgen) reveals that Eya proteins are expressed in murine immune cells, with Eya3 expressed ubiquitously across populations (Fig. 1A). Intriguingly, the level of Eya3 within macrophages is increased in mouse primary and metastatic breast tumors compared to normal breast[48], (Fig. 1B, Supp. Fig. 1A). We additionally analyzed single cell RNA-seq (scRNA-seq) data from human TNBC patients, and again found that Eya3 is expressed across TNBC infiltrating immune cell types, with higher expression found in innate immune populations such as NK cells, DCs, monocytes, and macrophages (Fig. 1C, Supp. Fig. 1B)[49]. Further, analysis of Eya3 expression in TNBC associated macrophages across a comprehensive study of eight patient cohorts reveals significantly increased macrophage Eya3 expression in grade 3 vs. grade 1 TNBC tumors (Fig. 1D)[50]. Taken together, these data suggest that Eya3 may play a prominent role in macrophages within the breast cancer microenvironment.

**Figure 1.**
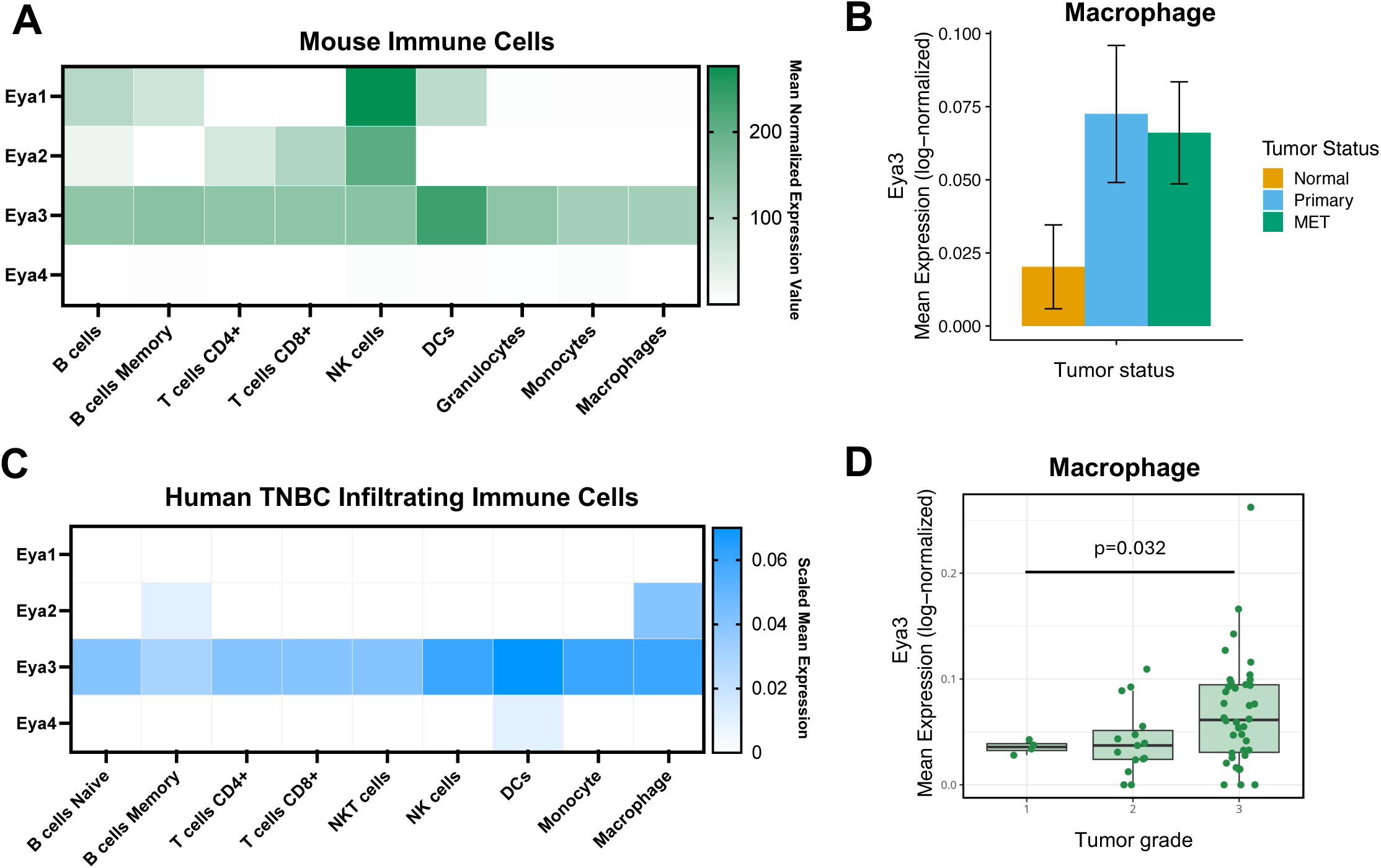
Eya3 is expressed in macrophages and correlates with pro-tumor phenotypes. A) Expression of Eya1-4 mRNA across mouse immune populations. Data sourced from the Immunological Genome Project ULI RNA-Seq database using the “My Geneset” tool (immgen.org). B) Macrophage Eya3 mRNA expression in murine normal breast, primary breast tumor, and lung metastasis from single cell RNA-seq study (GSE231915)[50]. C) Expression of EYA1-4 mRNA across human immune populations in TNBC. Data sourced from GSE176078[51] through the Broad Institute Single Cell Portal (https://singlecell.broadinstitute.org/singlecell). D) Tumor macrophage EYA3 expression (patient pseudobulk, n = 57 patients) in human grade 1-3 TNBC tumors across 8 studies[50].

### Loss of Eya3 globally alters gene expression, increasing migration and antigen processing in macrophage cell lines

To determine the role of Eya3 in macrophages, we first stably knocked down Eya3 using two independent short hairpin RNAs (shRNAs) in two macrophage cell lines: RAW264.7 and J774A.1 (Fig. 2A-B). We performed RNA-seq on RAW264.7 shSCR and shEYA3 KD cells and identified multiple differentially regulated pathways through gene set enrichment analysis that may impact macrophage function. We observed a negative net enrichment score (NES) with loss of Eya3 for “Myeloid Leukocyte Migration” (Supp. Fig. 2A), consistent with migration trends in cancer cells[19]. However, we found leading edge genes from this pathway to be predominantly composed of negatively enriched adhesion-related genes (Fig. 2C), and when examining alterations in a “matrix adhesion” gene set, we also observed overall negative enrichment (Supp. Fig. 2B). Furthermore, Eya3 loss led to a positive enrichment of multiple genes associated with actin polymerization, cytoskeletal remodeling, and lamellipodia formation (Fig. 2C). Taken together, these data strongly suggest that Eya3 may regulate macrophage motility (Fig. 2C), though the directionality of its effect was unclear.

**Figure 2.**
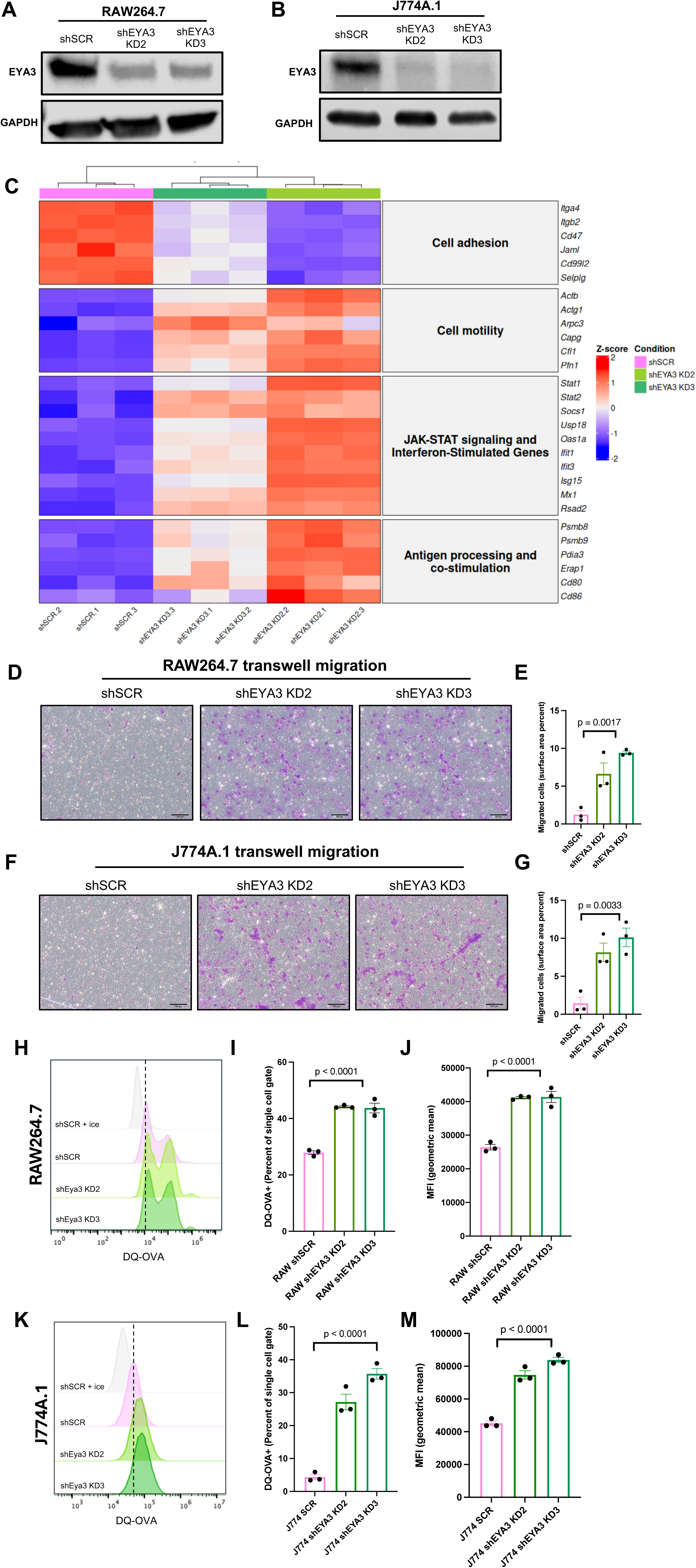
Loss of macrophage Eya3 alters gene expression of key pathways associated with macrophage function and leads to increases in migration and antigen processing. A, B) Western blot demonstrating knockdown of Eya3 in RAW264.7 and J774A.1 cells, respectively. C) RAW264.7 shSCR vs shEYA3 KD bulk RNA-seq summary heatmap of representative genes associated with relevant biological processes. RAW264.7 (D) and J774A.1 (F) 24hr transwell migration and associated quantification (E, G). Flow cytometry analysis of antigen processing in macrophage cell lines: H) RAW264.7 and K) J774A.1; representative histogram of DQ-OVA (FITC) signal. I, L) Percent DQ-OVA+ cells J, M) mean fluorescent intensity reported as geometric mean. Statistical analysis performed using one-way ANOVA. Experiments are representative of three independent biological experiments, which were each conducted with technical triplicates.

Gene expression analysis additionally demonstrated alterations to pathways that influence macrophage polarization states. We observed a negative NES with Eya3 KD for “TNFa Signaling via NFkB” (Supp. Fig. 2C), consistent with the known regulation of NFkB signaling by Eya in both *Drosophila* and in breast cancer[20, 51]. We also observed a striking enrichment with Eya3 KD in pathways associated with classically activated, “anti-tumor” macrophages, such as both type I and II interferon signaling, which impinges on JAK/STAT signaling (Fig. 2C, Supp. Fig. 2D-E). In addition, loss of Eya3 results in positive enrichment of “Antigen Processing and Presentation” gene sets (Supp. Fig. 2F-H). Macrophages and dendritic cells are professional antigen presenting cells (APC) that have been shown to acquire tumor associated neoantigens. These endocytosed antigens are processed by the APCs into short peptides for antigen presentation via MHC class I [52]. Because Eya3 has not previously been implicated in any specific immune functions, we further explored these findings via functional assays.

To determine how the alterations in gene expression influence key functions of macrophages, we first performed transwell migration assays with our control and Eya3 KD macrophage cell lines. In both the RAW264.7 and J774A.1 cells, loss of Eya3 significantly *increased* migration (Fig. 2D-G), contrary to the decreased migration observed with loss of Eya3 in breast cancer cells [19, 53]. These data suggest that the leading-edge genes that we identified in adhesion and actin-cytoskeleton may be dominant in driving the migratory phenotypes in macrophages.

We further examined antigen processing in control and Eya3 KD lines using DQ-OVA to evaluate proteolytic cleavage as a proxy for antigen processing. Intriguingly, we observe that loss of Eya3 in macrophage cell lines increases the fluorescence of DQ-OVA, an indication of increased antigen processing (Fig. 2H-M, Supp. Fig. 3). These data are consistent with the positive NES for antigen processing obtained from RNA-seq analysis (Fig. 2C, Supp. Fig. 2F-H), demonstrating that Eya3 function in macrophages intersects with proteolytic cleavage pathways. Taken together, these data strongly suggest a key role for Eya3 in macrophage function, though the results were limited to cell lines.

**Figure 3.**
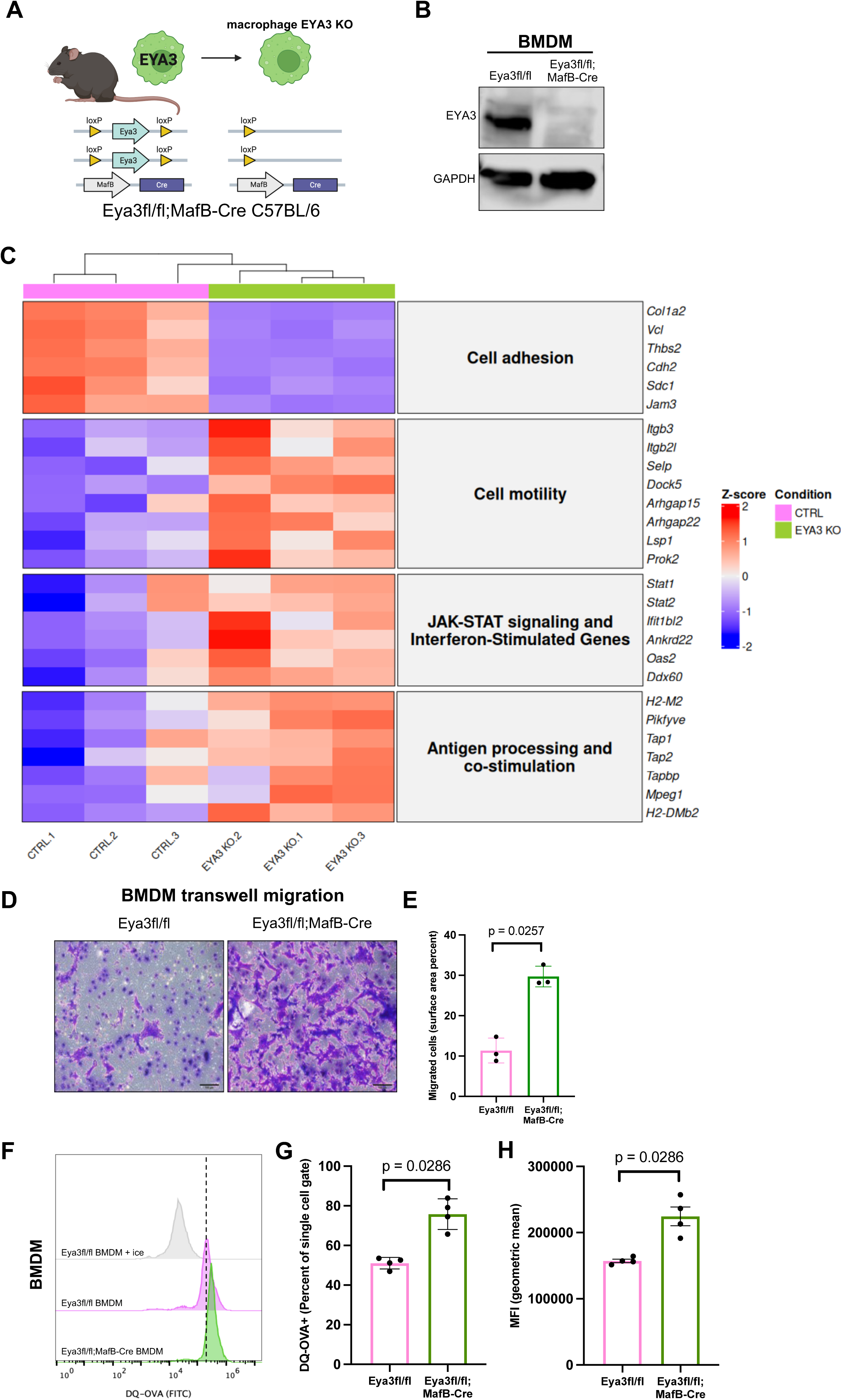
Macrophage-targeted Eya3 knockout reveals effects on gene expression, migration and antigen processing in primary macrophages, consistent with Eya3-deficient macrophage cell lines. A) Schematic representing generation of a macrophage Eya3 conditional knockout mouse through the Cre-lox system (MafB-Cre x Eya3-flox). B) Successful generation of macrophage Eya3 knockout mouse demonstrated through Western blot analysis examining Eya3 levels in bone marrow derived macrophages (BMDM). C) BMDM CTR (Eya3fl/fl) vs EYA3 KO (Eya3fl/fl;MafB-Cre) bulk RNA-seq summary heatmap of representative genes associated with relevant biological processes (GSEA plots in Supp. Fig.7). D) Eya3fl/fl and Eya3fl/fl;MafB-Cre BMDM 24hr transwell migration and E) quantification. F) BMDM antigen processing representative histogram G) Percent DQ-OVA+ BMDMs H) mean fluorescent intensity reported as geometric mean. Statistical analysis performed using unpaired t-test. All experiments are representative of at least three independent biological experiments, which were each conducted with at least technical triplicates.

### Generation and characterization of a macrophage Eya3 knockout mouse

Nearly two decades ago, a whole body Eya3 knockout (KO) mouse was developed. Eya3 loss led to minor, pleiotropic phenotypes in young mice, such as reduced muscle strength, decreased bone mineral content, decreased tidal volume of the lung at rest, and shorter body length [54]. In addition, a conditional Eya3 KO mouse was developed, which demonstrated a role for Eya3 in angiogenesis[26]. However, the immune system was not studied at baseline or during an immune challenge in either of these models. Thus, we developed a macrophage specific Eya3 conditional knockout mouse model to understand the role of Eya3 in this immune population.

To remove Eya3 specifically in macrophages, we crossed the previously generated Eya3-Flox mice[26], after backcrossing them onto the C57BL/6 background for eight generations (Supp. Fig. 4A), with MafB-Cre C57BL/6 mice (Fig. 3A, Supp. Fig 4A-B). MafB-Cre was chosen as the driver as it is the most specific Cre-driver for targeting macrophages within the immune system (Supp. Fig. 4C), though MafB is also required for the differentiation of osteoclasts, podocytes, and pancreatic beta cells[55–57]. In contrast, other Cre drivers that are used to generate mice with conditional knockout in macrophages, including LysM-Cre and Csf1r-Cre [56, 57], target additional cells in the immune system. For example, LysM-Cre targets dendritic cells, neutrophils, monocytes, and a population of lung epithelial cells, and Csf1r-Cre targets nearly all leukocytes[57]. Not only is MafB-Cre more specific to the immune system, but it is also highly expressed in both monocyte-derived macrophages, and most tissue resident macrophages[55], making it the ideal Cre-driver for our studies.

**Figure 4.**
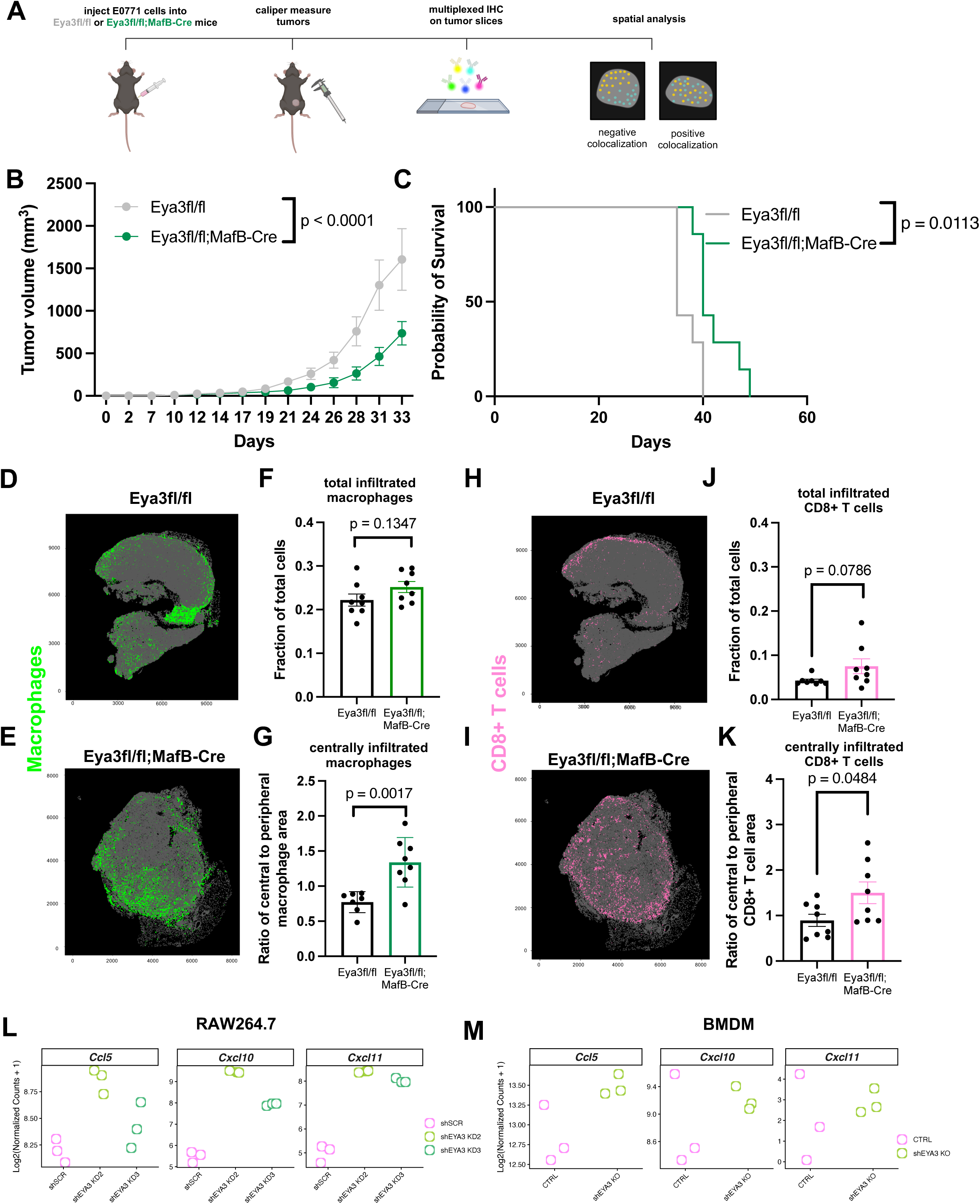
Macrophage specific Eya3 loss decreases primary TNBC growth in mice, accompanied by increased infiltration of macrophages and CD8+ T cells into tumors. A) Overview of experimental procedure. B) Primary tumor growth via caliper measurement after orthotopic injection of E0771 cells into Eya3fl/fl (n = 7) and Eya3fl/fl;MafB-Cre (n = 7) C57BL/6 mice (experiment 1). C) Survival to ethical endpoint of 1.5cm^3^ tumor volume for mice. D) CELESTA plots of all tumor infiltrating macrophages in Eya3 fl/fl vs. E) Eya3fl/fl;MafB-Cre tumor bearing mice. F) Total macrophages in Eya3 fl/fl and Eya3fl/fl;MafB-Cre tumors. G) Ratio of centrally localized to peripherally localized macrophages in Eya3 fl/fl and Eya3fl/fl;MafB-Cre tumors. H) CELESTA plots of tumor infiltrating CD8+ T cells in Eya3 fl/fl vs. I) Eya3fl/fl;MafB-Cre tumor bearing mice. J) Fraction of CD8+ T cells within Eya3 fl/fl and Eya3fl/fl;MafB-Cre tumors. K) Ratio of centrally localized to peripherally localized CD8+ T cells within Eya3 fl/fl and Eya3fl/fl;MafB-Cre. RNA expression dot plots of T cell recruiting chemokines from L) RAW264.7 and M) BMDM control and Eya3 KD/KO RNA-seq. Statistical analysis for panel B performed through 2-way ANOVA. Statistical analysis for panel C performed using log-rank test to compare survival. Statistical analysis for panels F-G, J-K performed using unpaired t-tests.

Successful knockout of Eya3 was confirmed by genotyping (Supp. Fig. 4D) and Western blot analysis of BMDMs (Fig. 3B). Flow cytometric characterization of macrophage populations at baseline (without an immune challenge) from the blood, lung, spleen, lymph node, and mammary gland from control Eya3fl/fl, vs. macrophage specific Eya3 knockout Eya3fl/fl;MafB-Cre mice (Supp. Fig. 4E-I, Supp. Fig. 5) revealed only modest changes in macrophage quantity in some compartments, with the only significant decrease found in spleen yolk sac derived macrophages (Supp. Fig. 4G). When examining monocytes, dendritic cells, CD4+ T cells, CD8+ T cells, NK cells, B cells, and neutrophils across the tissues analyzed, the only significant change observed was a decrease in blood NK cells (Supp. Fig. 5, Supp. Fig. 6E shows the gating strategy). These data demonstrate that the immune system of macrophage Eya3 KO mice remains largely intact at baseline.

**Figure 5.**
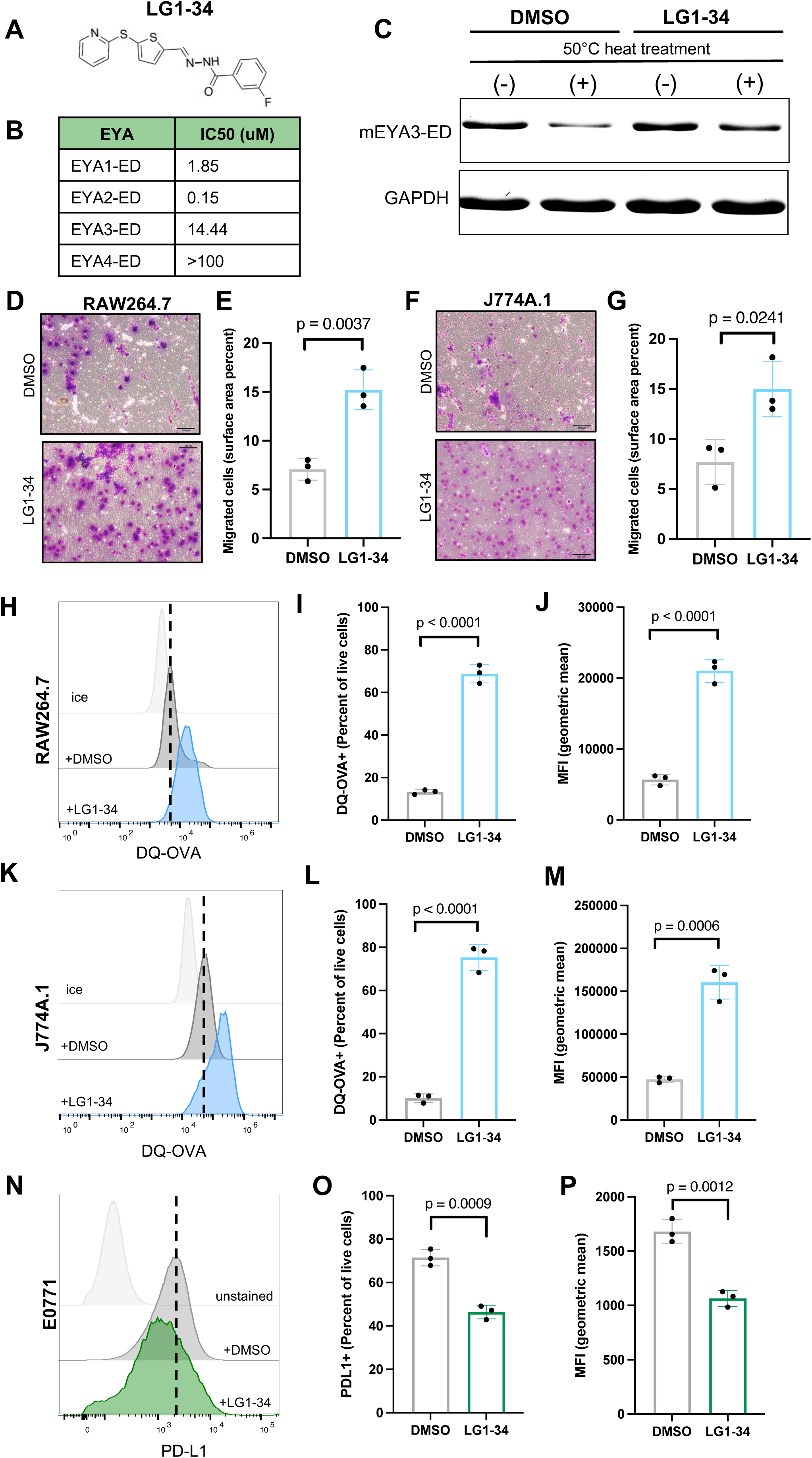
LG1-34, a novel allosteric small molecule Eya1-3 tyrosine phosphatase inhibitor, phenocopies genetic loss of Eya3 in macrophages. A) Structure of LG1-1. 34. B) LG1-34 IC50s for purified Eya1-4 protein using an *in vitro* phosphatase assay. C) Thermal stability assay (TSA) performed using purified Eya3 protein (Eya domain (ED) only) incubated with DMSO or 20uM LG1-34, at 50°C. Supernatant analyzed on SDS-PAGE and stained by SYPRO Ruby. Spiked in GAPDH was used as a loading control. 24hr transwell migration assay performed on RAW264.7 (D) or J774A.1 (F) cells treated with DMSO or 2uM LG1-34. Quantitation shown in E, G respectively. H-J) Flow cytometric analysis of antigen processing in RAW264.7 (H) or J774A.1 (K) cells treated with 2uM LG1-34 for 72hr, then incubated with DQ-OVA. Representative histograms (H,K), percent DQ-OVA+ cells (I, L), and geometric mean fluorescent intensity (J, M) are shown. Flow cytometric analysis of PD-L1 expression on E0771 cells after treatment with 2uM LG1-34 for 72h: Representative histogram (N), percent PD-L1+ cells (O), and geometric mean fluorescent intensity (P) of PD-L1 are shown. Statistical analysis performed using unpaired t-tests. Experiments are representative of three biological experiments, which were each conducted with technical triplicates.

**Figure 6.**
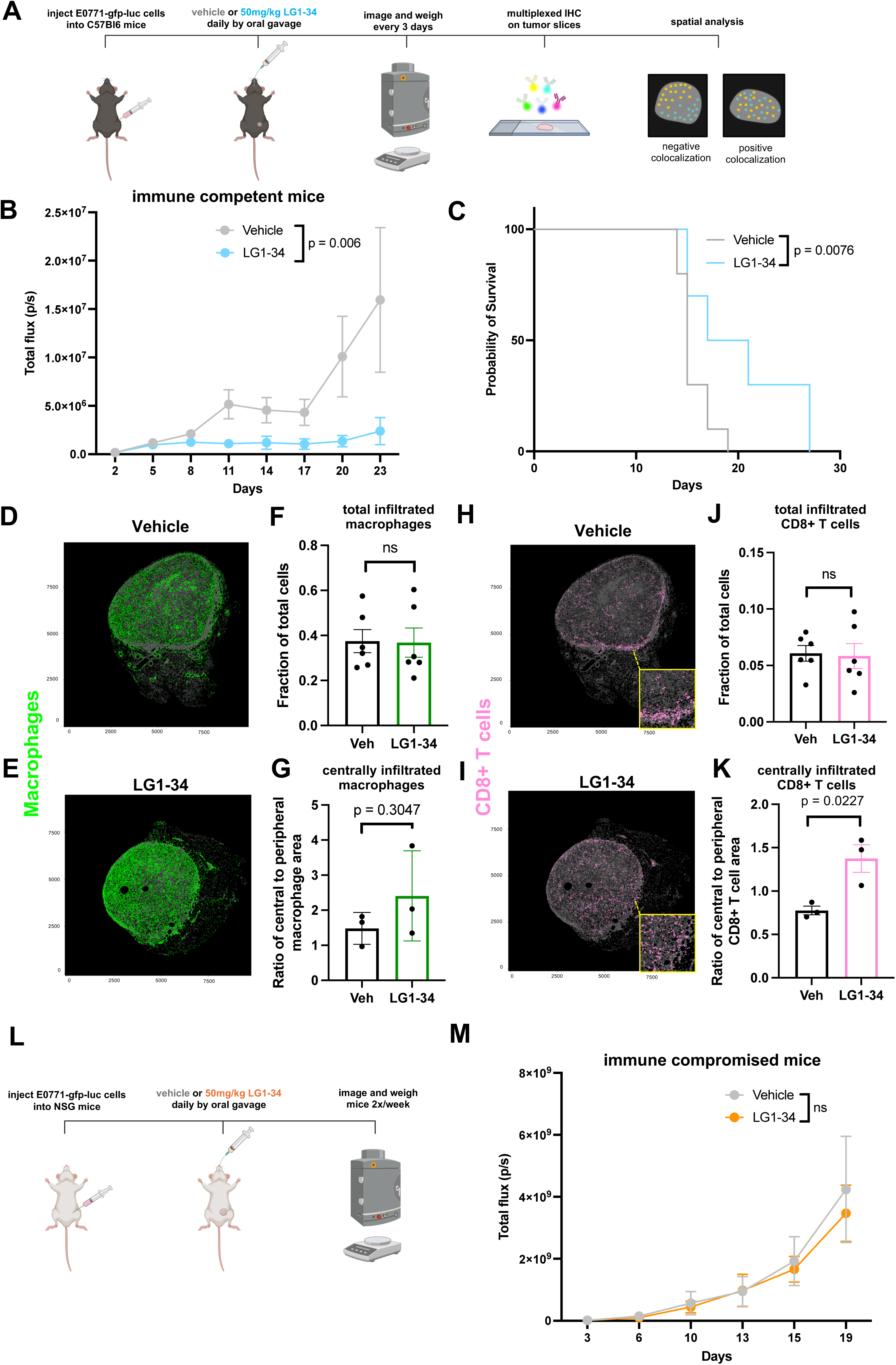
Inhibition of Eya’s tyrosine phosphatase activity with LG1-34 suppresses TNBC tumor growth by enhancing immune-mediated tumor elimination. A) Overview of immune competent LG1-34 tumor study. B) Primary tumor growth via bioluminescent imaging after orthotopic injection of luciferase-tagged E0771 cells into vehicle (n = 10, 6 tumors analyzed by IHC and spatial analysis) and LG1-34 (n = 10, 6 tumors analyzed by IHC and spatial analysis; 4 tumors had complete response to treatment) treated C57BL/6 mice. Mice were treated with 50mg/kg or an equivalent volume of DMSO daily by oral gavage. Data shown for *in vivo* experiment #1, growth curves from experiment #2 shown in Suppl. Fig. 9B. C) Survival to ethical endpoint for mice (from LG1-34 *in vivo* experiment #2). D) CELESTA plots of all tumor infiltrating macrophages in vehicle treated and E) LG1-34 treated mice. F) Fraction of macrophages within tumors. G) Ratio of centrally localized to peripherally localized macrophages from n = 3 tumors per group, size-matched small tumors (see Supp. Fig. 15A). H) CELESTA plots of tumor infiltrating CD8+ T cells in vehicle treated and I) LG1-34 treated mice. J) Fraction of CD8+ T cells within tumors. K) Ratio of centrally localized to peripherally localized CD8+ T cells from n = 3 tumors per group, size-matched small tumors. L) Overview of immune compromised LG1-34 tumor study. M) Primary tumor growth via bioluminescent imaging in NSG mice (n = 10 vehicle treated, n = 10 LG1-34 treated mice). Statistical analysis for panels B and M was performed through 2-way ANOVA. Statistical analysis for panel C was performed using log-rank test to compare survival. Statistical analysis for panels F-G and J-K was performed using unpaired t-tests.

### Eya3 is a critical regulator of primary macrophage gene expression, migration, and antigen processing

To interrogate the function of Eya3 in primary macrophages, we isolated bone marrow from Eya3fl/fl or Eya3fl/fl;MafB-Cre mice and cultured the cells *ex vivo* to differentiate into bone marrow derived macrophages. As observed in the RAW264.7 Eya3 KD macrophage cell line, bulk RNA-seq analysis of control and Eya3 KO BMDMs identified similar enrichment patterns among gene sets associated with cell motility, macrophage activation, and antigen processing and presentation (Fig. 3C, Supp. Fig. 7). Consistent with our results *in vitro*, we observed significant increases in transwell migration (Fig. 3D-E) as well as in antigen processing (Fig. 3F-H), with Eya3 KO in BMDMs.

**Figure 7.**
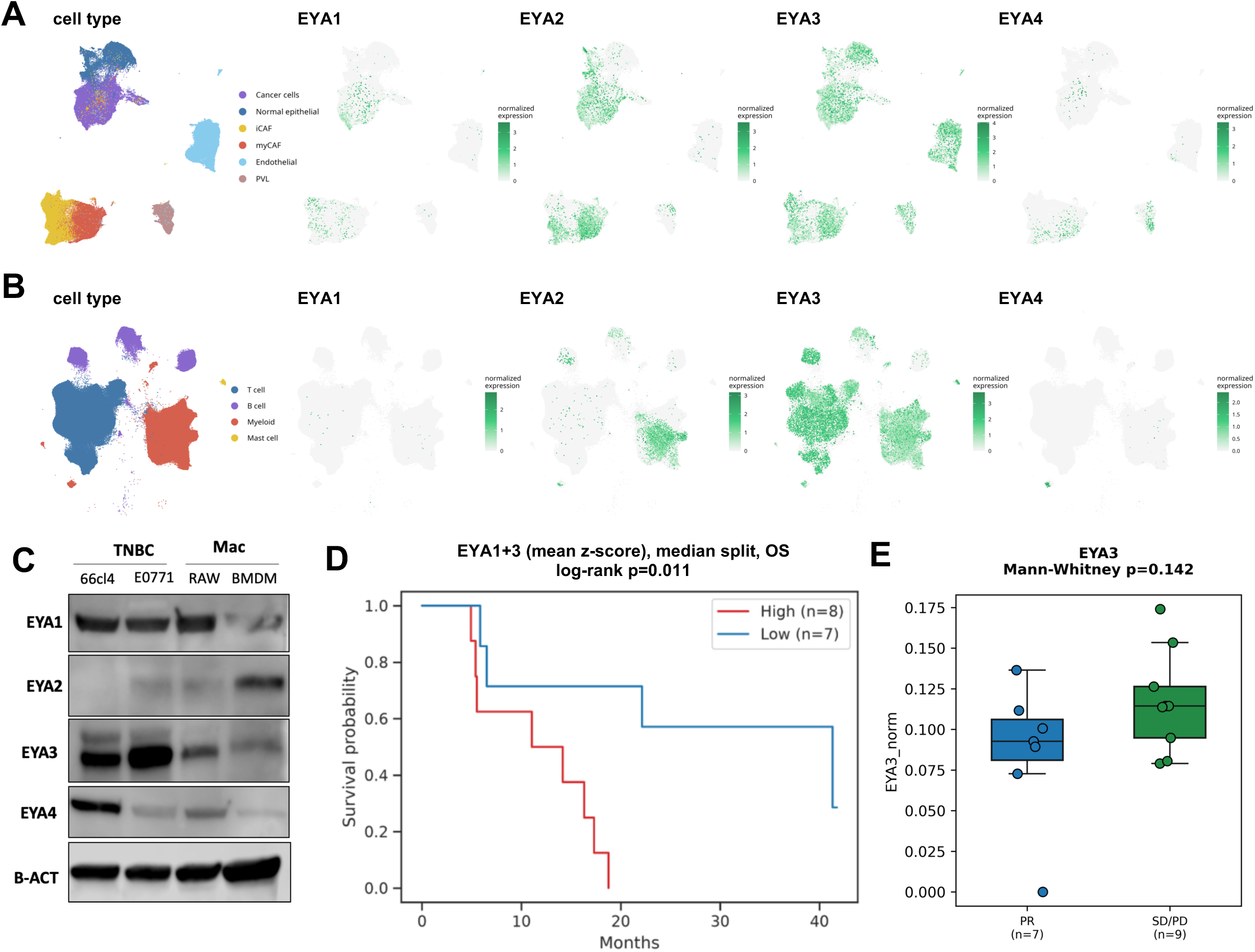
High Eya expression correlates with worse overall survival and treatment response in TNBC following immunotherapy. A) UMAPs of Eya1-4 RNA expression in cancer, normal epithelial, and stromal cells (iCAF, myCAF, endothelial, and PVL (perivascular-like cells)) and B) immune cells (T-cells, B-cells, Myeloid cells, and Mast cells) from TNBC patients following pembrolizumab (anti-PD-1) and radiation therapy (GSE246613). C) Western blot analysis of Eya1-4 protein expression in TNBC cell lines 66cl4 and E0771, macrophage cell line RAW264.7, and bone marrow derived macrophages (BMDM) from whole cell lysates. D) Overall survival of TNBC patients with high and low Eya1 + 3 expression following pembrolizumab (anti-PD-1) + carboplatin/nab-paclitaxel treatment (GSE241876). E) Tumor macrophage Eya3 expression in TNBC patients with partial response (PR), stable disease (SD), or progressive disease (PD) following paclitaxel plus atezolizumab (anti-PD-L1) treatment (GSE169246).

We further evaluated whether the difference in antigen processing would affect presentation of antigen to CD8+ T cells, as this is a critical communication step between the innate and adaptive immune responses, and can influence tumor growth. To assess antigen presentation, we isolated CD8+ T cells from OT-1 mice, a T cell receptor transgenic mouse in which all T cell receptors recognize the SIINFEKL peptide in the context of the major histocompatibility complex I molecule H-2K^b^. We co-cultured violet proliferation dye (VPD)-labeled OT-1 CD8+ T cells with Eya3fl/fl or Eya3fl/fl;MafB-Cre BMDMs that were incubated with OVA protein (requires processing prior to presentation), or SIINFEKL peptide (does not require processing). After 3 days of co-culture, we measured OT-1 proliferation by dilution of the proliferation dye (Supp. Fig. 8A). We observed a modest, but significant, increase in the proliferation index, a measure of the average number of times the responding T cells divided, in OT-1s when either OVA or SIINFEKL was given to the EYA3 KO BMDMs for SIINFEKL presentation (Supp. Fig. 8B-D). Together, our findings in macrophage cell lines and primary macrophages indicate that loss of Eya3 enhances functions that are associated with classically activated macrophages, and could thus play a critical role in regulating tumor growth within the TIME.

### Loss of macrophage Eya3 decreases primary TNBC growth in mice, and leads to increased infiltration of macrophages and CD8+ T cells into tumors

To determine whether the observed regulation of macrophage function by Eya3 may influence key *in vivo* functions of macrophages, such as their effects on tumor growth, we injected E0771 cells into the mammary fat pad of syngeneic control Eya3fl/fl and Eya3 knockout Eya3fl/fl;MafB-Cre C57BL/6 mice and assessed tumor growth by caliper measurement (Fig. 4A). Intriguingly, loss of Eya3 specifically in the macrophage compartment (and not in the tumor cells) led to significantly decreased tumor growth (Fig. 4B, Supp. Fig. 9A), and mice with macrophage Eya3 KO show improved survival when taken to ethical endpoint (Fig. 4C). Furthermore, the MafB-Cre control mice develop tumors statistically identical in size to Eya3fl/fl control mice, suggesting no effect of Cre itself on the TIME (Supp. Fig. 9B-C), and thus experiments throughout this work were carried out using Eya3fl/fl mice as the control for mice with Eya3-deficient macrophages.

To understand how Eya3 KO in macrophages influences tumor growth, we examined the presence of various immune populations in the tumors using both flow cytometry and multiplexed fluorescent immunohistochemistry (mIHC). Flow cytometry analysis did not reveal statistically significant changes in total macrophage numbers in tumors from Eya3fl/fl or Eya3fl/fl;MafB-Cre mice at ethical endpoint (Supp. Fig. 10A-B). Similarly, we observed no significant differences by flow cytometry in the total numbers of other immune cell types within tumors, including CD8+ T cells (Supp. Fig.5, Supp. Fig. 10C-F).

To assess whether immune cells exhibit changes in their spatial distribution within tumors from Eya3fl/fl and Eya3fl/fl;MafB-Cre mice, we analyzed multiplexed IHC images of tumors taken earlier in disease progression (Supp. Fig. 11A). CELESTA[58] was used to identify cell populations (Fig. 4D-K, Supp. Tables 1-2, Supp. Fig. 11). While we again did not observe a difference in total infiltration of any immune population identified through our 7-marker panel (Supp. Fig. 11B, E), we did observe a significant difference in the ratio of both macrophages (all sub-populations combined), and CD8+ T cells that are centrally infiltrated vs. peripherally localized in tumors from Eya3fl/fl;MafB-Cre mice when compared to their control counterparts (Fig. 4G, K). Further, we generated colocation quotients (CLQs)[59] between all cell population pairs (Supplemental Excel file 1), with a positive normalized CLQ indicating colocation or occupation of the same spatial niche, and a negative normalized CLQ indicating segregation or occupation of distinct spatial niches[60]. Through this quantitative analysis, we identified an additional metric for the central infiltration of CD8+ T cells. CD8+ T cells in tumors from Eya3fl/fl;MafB-Cre mice were significantly less colocated with a sub-population of peripherally-localized macrophages that express F480+, CD80+, CD163+, and EGFR+ (Supp. Fig. 11C-D), when compared to CD8+ T cells in control Eya3fl/fl mice (Supp. Fig. 11F-H), resulting in a greater negative normalized CLQ (Supp. Fig. 11H). These data demonstrate that loss of Eya3 results in alterations to the spatial organization between macrophages and CD8+ T cells, and greater infiltration of CD8+ T cells into the tumor.

Once CD8+ T cells are primed to carry out cytotoxic effector functions, they must home back to the tumor to mount an anti-tumor response, and they do so in part by following chemokine gradients[61, 62]. From our RAW264.7 and BMDM Eya3 KD/KO RNA-seq analysis, we identified T-cell recruitment gene modules that include chemokines such as CCL5, CXCL10, and CXCL11 that have increased expression with loss of macrophage Eya3 (Fig. 4L-M). Collectively, these findings suggest that expression of Eya3 within macrophages leads to enhanced tumor growth through multiple mechanisms whereby Eya3 augments pro-tumor macrophage functions, suppressing chemokine gradients and central CD8+ T-cell infiltration. Loss of Eya3 reverses these effects, leading to macrophages that are more tumor suppressive in function, and importantly, to enhanced CD8+ T-cell infiltration, thus effectively altering the TNBC TIME towards a “hotter” immune phenotype.

### LG1-34, a novel allosteric Eya tyrosine phosphatase inhibitor, suppresses TNBC tumor growth by enhancing immune-mediated tumor elimination

As loss of Eya3 in both the macrophage (Fig. 4B) and tumor[19] compartments leads to decreased tumor growth and increased CD8+ T cell infiltration, we examined whether targeted inhibition of Eya3 may lead to immune-dependent tumor elimination. To this end, we modified a previously developed allosteric inhibitor of the Eya2 tyrosine phosphatase to also inhibit the Eya3 tyrosine phosphatase. Eya proteins contain several functional domains, including an N-terminal transactivation domain, which also carries out PP2A-associated serine/threonine phosphatase activity, and a C-terminal EYA domain, which interacts with the SIX family of homeoproteins, and also harbors intrinsic tyrosine phosphatase activity[63]. We have previously shown that inhibition of the Eya2 tyrosine phosphatase activity can impinge on its associated PP2A threonine phosphatase activity[64], thereby decreasing Myc levels. As we have also shown that Myc is critical for Eya3 to inhibit the CD8+ T-cell response[19], we reasoned that inhibition of the Eya3 tyrosine phosphatase activity may impair the immune suppressive functions of Eya3 in both the macrophage and the tumor compartment. Through medicinal chemistry efforts, we developed a new allosteric inhibitor, LG1-34[46], that can target EYAs 1 and 3, albeit with lower potency (higher IC50s, as seen in *in vitro* phosphatase assays using purified Eya proteins) than observed for Eya2 (Fig. 5A-B). Because Eya3 is the dominant Eya in our models, we performed thermal stability assays (TSA) with purified Eya3 to demonstrate that our novel allosteric inhibitor can stabilize Eya3 against heat denaturation, and thus binds Eya3 directly (Fig. 5C). We then treated macrophage cell lines with LG1-34 [46], and determined that similar to Eya3 KD or KO, inhibition of the Eya tyrosine phosphatase activity increased migration (Fig. 5D-G) and processing in these contexts (Fig. 5H-M). Since our previous data demonstrated Eya3 functions within tumor cells themselves, we also treated E0771 cells with LG1-34, and found that our tyrosine phosphatase inhibitor leads to decreased surface expression of PD-L1 on the tumor cells (Fig. 5N-P, Supp. Fig.12 shows gating strategy), mirroring the effect of Eya3 KD in TNBC that was linked to an improved CD8+ T cell response[19]. Taken together, our data indicate that LG1-34 may inhibit tumor progression by enhancing CD8+ T cell responses via targeting Eya family tyrosine phosphatase activity in both tumor cells and macrophages.

Our *in vitro* data suggest that inhibiting Eya3 in the tumor and macrophage compartments may have an additive effect on inhibition of tumor growth by enhancing the immune response. To test whether pharmacologic inhibition of Eya3 may be a powerful means to restore immune-mediated tumor elimination, we injected the mammary fat pads of immune competent C57BL/6 mice with luciferase-tagged E0771 TNBC cells, after which we treated mice with vehicle or 50 mg/kg LG1-34 daily by oral gavage (Fig. 6A). Bioluminescent imaging over the course of three weeks revealed dramatic inhibition of primary tumor growth in mice treated with LG1-34 (Fig. 6B, Supp. Fig. 13A-B). Furthermore, in a second experiment in which the tumors were taken to a humane endpoint, we found that LG1-34 treated mice had a significant survival advantage (Fig. 6C, growth curve for this experiment shown in Supp. Fig. 13C). Importantly, the significant effect on tumor growth was seen without any changes in body weight, demonstrating that the compound was well-tolerated by the mice (Supp. Fig. 13D-E).

To examine the TIME in LG1-34 treated tumors, we euthanized mice at day 23 from the first *in vivo* experiment, and performed mIHC followed by CELESTA (Fig. 6A). As seen in our macrophage specific Eya3 KO experiment, we did not observe significant differences in the quantity of tumor associated immune cells between vehicle or LG1-34 groups (Fig. 6F,J, Supp. Table 3, Supp. Fig. 14A-B). Importantly, however, we did observe a trend towards increased central localization of macrophages (Fig. 6G), and a statistically significant increase in central localization of CD8+ T cells with LG1-34 treatment, a key shared spatial feature between similarly sized tumors (∼200mm^3^) from macrophage Eya3 knockout mice (Fig. 4I, K) and LG1-34 treated mice (Fig. 6I, K). These data, together with our *in vitro* and *ex vivo* data, suggest that LG1-34 treatment inhibits tumor growth in part by improving the CD8+ T cell response, likely through effects both on the tumor and macrophage compartments.

While LG1-34 treatment largely phenocopies genetic loss of Eya3, a prominent, unique spatial feature of tumors from LG1-34 treated mice vs. macrophage Eya3 knockout mice, identified through CLQ comparison between populations and treatment groups (Supplemental Excel file 2), is increased colocation of CD3+ T cells (CD3+, CD8-), which are likely CD4+ T cells, and “general macrophages” (F480+, CD80-, CD163-, EGFR-) with LG1-34 treatment (Supp. Fig. 14C-E). This interaction could indicate additional macrophage – T cell signaling that influences the TIME.

To test whether the anti-tumor effect of LG1-34 in TNBC-bearing mice is driven through an immune response, whether acting on tumor cells or immune cells, we repeated this study in immune-compromised NSG mice. As anticipated, we observed no difference in tumor growth between LG1-34 treated and vehicle groups in mice lacking an intact immune system (Fig. 6L-M, Supp. Fig. 15). Collectively, these findings establish Eya tyrosine phosphatase inhibition as a novel strategy to remodel the TIME and augment anti-tumor immunity, providing pre-clinical rationale for targeting Eya in TNBC.

### High Eya3 expression in macrophages correlates with decreased response to anti-PD1 treatment in TNBC patients

To determine how macrophage Eya3 expression may affect response to immunotherapy in TNBC patients, we analyzed Eya expression within the TNBC TIME following anti-PD-1 or anti-PD-L1 therapy from publicly available datasets. Analysis of scRNA-seq from a clinical study in which patients were treated with pembrolizumab and radiation therapy[65], demonstrates that Eya3 is the most abundantly expressed Eya across the tumor and immune compartments, with significant expression in the macrophage population (Fig. 7A-B). Western blot analysis of TNBC, macrophage cell lines, and BMDMs revealed a similar pattern, whereby Eya3 protein is highly expressed across both the tumor and macrophage cells, followed by Eya1 (Fig. 7C), which the scRNA-Seq data demonstrates is primarily expressed by cancer cells (Fig. 7A). Because our small molecule Eya inhibitor targets Eya1-3, and because all three of the Eyas are expressed in the tumor and/or immune compartments (Figs. 7A-B), we examined overall survival (OS) with high/low stratified Eya expression via median split following pembrolizumab (anti-PD-1) and carboplatin/nab-paclitaxel treatment in a phase II trial[66], and identified a significant advantage in survival probability when stratifying by high and low Eya1 and 3 expression together (Fig. 7D), with significance for EYA1 alone and a trend towards significance for Eya3 alone (Supp. Fig. 16), though it should be noted that the patient number in this study was small. Importantly, in pre- and post-treatment scRNA-seq profiles from TNBC patients who underwent atezolizumab (anti-PD-L1) plus paclitaxel treatment [67], elevated Eya3 expression in tumor-associated macrophages showed a trend toward enrichment in patients with stable disease (SD) or progressive disease (PD) relative to responding patients (Fig. 7E; p = 0.174). This pattern was specific to macrophages and was not observed in monocytes (Supp. Fig. 17A), and is consistent with a potential association between macrophage EYA3 and therapeutic resistance. Together, these findings suggest that elevated Eya3 expression, particularly in macrophages, is associated with poorer outcomes and reduced response to immunotherapy in TNBC, supporting therapeutic targeting of Eya3 as a potential strategy to enhance treatment response.

## Discussion

Discovering novel immunotherapeutic targets in TNBC is critical to improving patient treatment outcomes. In this study, we identify the developmental protein Eya3 as a previously unrecognized regulator of macrophage function within the TIME and demonstrate that its inhibition enhances anti-tumor immunity in TNBC. While Eya3 has been implicated in tumor cell intrinsic processes that promote tumor progression, as well as tumor suppression of the immune response, our findings expand its role to the immune compartment, revealing that Eya3 can act across both tumor and immune cells to coordinately suppress cytotoxic immune responses. This dual-compartment activity positions Eya3 as a unique player in the TIME and suggests that inhibiting this shared signaling program may offer a therapeutic advantage over approaches that target a singular cellular compartment.

Because of their plasticity and abundance within solid tumors, macrophages are an intriguing target for cancer immunotherapy[43, 68, 69]. While tumor-associated macrophages (TAMs) have historically been thought of as a negative prognostic marker for several solid tumor types[70], it has since been recognized that nuances in macrophage function, spatial localization within a tumor, and cell-cell interactions dictate the “pro-tumor” or “anti-tumor” effects of this immune population far more than abundance alone[8]. Therefore, it is critical that we continue to uncover means to promote anti-tumor functions of macrophages as a method of limiting tumor growth[8].

Our results align with the paradigm shift that macrophage function outweighs the effect of macrophage abundance alone, or even polarization state, within a tumor. While it is true that macrophages are extraordinarily plastic, they do not have to express canonical “M1” or “M2” markers to have anti- or pro-tumor functions, respectively. Further, they can express a combination of both. In the context of our experiments, we did not observe a significant change in the fraction of M1 macrophages (CD80+), or M2 macrophages (CD163+) (Supp. Fig. 11D, 14B), nor did we observe a significant difference in the number of total macrophages when comparing macrophage Eya3 KO mice to control mice. Instead, we identify a difference in the spatial organization of macrophages and CD8+ T cells within the tumor (increased central infiltration), as well as altered spatial relationships between specific T cell and macrophage populations, with loss or inhibition of macrophage Eya3. For example, increased interaction between CD4+ T cells (the likely identity of CD3+/CD8-T cells) and unpolarized macrophages in tumors of LG1-34 treated mice (Supp. Fig. 14C-E) could contribute to an anti-tumor immune response, as Th1 CD4+ T cells can support anti-tumor functions of macrophages, and macrophages can activate or re-activate CD4+ T cells [71, 72]. Therefore, our data suggest that alterations in macrophage function and spatial organization driven by Eya3, rather than the abundance or binary polarization state, drive the differences in tumor growth.

Unique features in tumors of LG1-34 treated mice compared to tumors from mice with macrophage Eya3 KO or TNBC Eya3 KD may be due to inhibition of Eya3 across multiple additional compartments (not only cancer cells and macrophage, but also in other immune cells within the TIME). Alternatively, these additional effects could occur due to inhibition of multiple Eyas within the TIME, as LG1-34 targets Eya1-3. Finally, variance could be observed due to differences between inhibition of tyrosine phosphatase activity specifically, rather than full depletion of the protein, which would remove multiple Eya activities including its Tyr and Ser/Thr phosphatase activities, as well as its transcriptional activity.

Boosting antigen processing and presentation may be an effective strategy for improving cytotoxic T-cell responses in heterogenous tumors with elevated levels of neoantigen, such as TNBC. There is precedent for neoantigen-based therapies, and a recent individualized mRNA vaccine for adjuvant TNBC has proven effective in a number of patients[73]. Other neoantigen-based therapies include neoantigen-loaded dendritic cell vaccines[74], and APC-targeted liposome-neoantigen vaccines[75]. While our compound, LG1-34, does not amplify neoantigen exogenously and re-introduce it as these vaccine strategies do, it boosts the ability of APCs *in situ,* such as macrophages, to generate an anti-tumor immune environment. In addition, by altering PD-L1 levels on the tumor cells, it alters the immune response both through effects on tumor and macrophage populations. Personalized approaches to cancer therapy, such as the TNBC mRNA vaccine[73], offer incredible therapeutic potential, but have an inherent lag in time-to-treatment based on the time required to acquire patient material, engineer the vaccine to reflect the neoantigens of the specific patient, and then begin treatment. It is thus essential to develop complementary therapeutic strategies to enhance the immune response to tumors, such as could be achieved through targeting Eya3.

Previous work showed that Eya3 promotes surface PD-L1 expression by stabilizing Myc, which in turn suppresses the cytotoxic T cell response[19]. Here, we show that use of a novel, systemic allosteric Eya tyrosine phosphatase inhibitor, LG1-34, decreases tumor growth, and likely does so through multiple mechanisms that converge on altering CD8+ T-cell responses to the tumor. We show that the decreased tumor growth observed with LG1-34 treatment is likely in part via its inhibition of Eya3 in macrophages. LG1-34 may enhance tumor elimination by promoting macrophage migration and antigen processing while reducing PD-L1 expression on tumor cells. Together, these effects may increase CD8+ T-cell localization within tumors, thereby enhancing anti-tumor immune responses.

In summary, these findings support a model in which Eya3 functions as a shared regulator of tumor and immune cell behavior, coordinating programs that jointly limit anti-tumor immunity. By constraining chemokine expression, migration, antigen processing in macrophages, Eya3 dampens a critical communication axis between innate and adaptive immune populations, which ultimately restricts the tumor-infiltrating cytotoxic T cell response. Importantly, inhibition of macrophage Eya3, specifically its tyrosine phosphatase activity, reverses the immune suppressive functions of Eya3 and significantly decreases primary tumor burden in mice. Given the limited treatment options for patients with TNBC, this work acts as an exciting, proof-of-concept study for Eya3 inhibition as a targeted, yet, cross-compartmental, treatment for TNBC. As this is, to our knowledge, the first investigation into the role of Eya3 in macrophages, future work defining the molecular substrates of Eya3 in macrophages, as well as testing Eya inhibition in combination with existing immunotherapies, will be critical for improved mechanistic understanding and movement of our small molecule inhibitor towards clinical use. Collectively, this work suggests that disrupting developmental programs like Eya3 may be a powerful therapeutic strategy to limit tumor growth and restore anti-tumor immunity in aggressive, immunogenic cancers such as TNBC.

## Methods

### Analysis of immune Eya expression in publicly available datasets

Analysis of Eya1-4 RNA expression across immune cell types in mice was performed using the Immunological Genome Project’s “My Geneset” tool on their ULI RNA-seq dataset (https://www.immgen.org/). To analyze macrophage Eya3 expression across normal, primary, and metastatic breast tissue, Eya3 expression in macrophages was calculated from single-cell RNA-seq from the study “Spatial and Temporal Mapping of Breast Cancer Lung Metastases Identify TREM2 Macrophages as Regulators of the Metastatic Boundary” (GSE231915)[48]. Log-normalized Eya3 expression was extracted from the macrophage subset using Seurat. Mean expression +/- standard error was calculated across cells grouped by tumor status (Normal, Primary, Metastatic). Analysis of Eya1-4 mRNA expression across immune cell types in human TNBC was performed through the Broad Institute’s Single Cell Portal (https://singlecell.broadinstitute.org/single_cell). Data was sourced from TNBC patients from the study “A single-cell and spatially resolved atlas of human breast cancers” (GSE176078)[49]. TIMER2.0 (https://compbio.cn/timer2/), an online tool for assessing immune infiltrates across cancer types, was used to identify M2 macrophage correlation with Eya3 expression in the TCGA breast cancer dataset via CIBERSORT analysis[76].

### Cell lines and culture conditions

Murine macrophage cell line RAW264.7 was purchased from ATCC (TIB-71, Manassas, VA) and cultured in Dulbecco’s modified Eagle’s medium (DMEM) high-glucose media supplemented with 10% fetal bovine serum (FBS), 0.5% penicillin/streptomycin (pen/strep), and 1% sodium pyruvate, referred to moving forward as “macrophage media”. Murine macrophage cell line J774A.1 was obtained as a gift from T. Lyons and cultured in macrophage media. Murine TNBC cell lines E0771 and E0771-gfp-luc were obtained as a gift from D. Cittelly and cultured in Roswell Park Memorial Institute (RPMI) media supplemented with 10% FBS and 1% pen/strep. All cell lines were grown at 37°C in 5% CO_2_, checked for mycoplasma every 6 months, and replaced once near or at passage 20.

### Generation of Eya3 knockdown cell lines

Eya3 was knocked down in macrophage cell lines using two different murine Eya3-targeting short hairpin RNAs (shRNA): shEya3 KD2-TRCN0000029858 and shEya3 KD3-TRCN0000029855 (Dharmacon, Lafayette, CO), alongside a control scramble shRNA (Addgene plasmid 1864, Cambridge, MA). The shRNAs were lentivirally introduced into cells according to the PLKO.1 manufacturing protocol (Addgene, Cambridge, MA), and pooled Eya3 KD populations were generated through puromycin selection (5ug/mL for RAW264.7 cells and 1ug/mL for J774A.1 cells).

### Transwell migration assays

Macrophages were plated at a concentration of 200,000 cells/200ul serum-free media per transwell insert with 8um pores (BD Falcon, 353097), with triplicate inserts plated per group. Transwell inserts were placed within 24-well companion plates (Corning, 353504), each well containing 800uL of full serum macrophage media and incubated at 37°C in 5% CO_2_ for 24hr. For migration assays with LG1-34, 2uM treatment began at the start of this 24hr period. After incubation, the bottom of each insert was fixed in 10% buffered formalin for 10m at RT. After fixation, the bottom of each insert was stained with 0.1% crystal violet solution for 45m at RT, then washed with ddH20 to remove excess stain, and any cells remaining within the transwell (not cells that migrated through to the underside of the membrane) were removed with a cotton swab. The transwell inserts were allowed to dry at RT for 24hr, then imaged on an Olympus CKX41 microscope at 10X and analyzed using ImageJ.

### Western blot analysis

Whole cell protein lysates (WCL) were generated by lysis of pelleted cells in radioimmunoprecipitation assay (RIPA) buffer supplemented with protease inhibitor (Thermo Scientific, a37989) and PhosSTOP phosphatase inhibitor (Roche, 04906845001) and further lysed through sonication. Protein was quantified by Lowry assay, then 50ug of protein was electrophoresed on a gradient acrylamide gel and transferred to a polyvinylidene difluoride (PVDF) membrane. Membranes were blocked for 1hr in 5% bovine serum albumin (BSA) in tris-buffered saline with Tween 20 (TBST), incubated in primary antibody (diluted in 5% BSA in TBST) overnight at 4°C, then incubated in secondary antibody (diluted in 5% non-fat dry milk in TBST) for 1hr at room temperature before imaging. Blots were developed with SuperSignal West Pico Chemiluminescent Substrate (Thermo Fisher Scientific, 34080) and/or SuperSignal West Femto Chemiluminscent Substrate (Thermo Fisher Scientific, 34096) and imaged on a Licor Odyssey FC instrument. Relevant antibodies used can be found in Supplemental Table 2.

### Generation of macrophage Eya3 conditional knockout mouse

A pair of homozygous Eya3-flox mice was obtained from R. Hegde[26]. These mice were on a mixed genetic background, predominantly Agouti, and containing some amount of C57BL/6. We backcrossed these mice onto a C57BL/6 genetic background for 8 generations, at which point E0771 TNBC cells, syngeneic to C57BL/6, were not rejected when orthotopically injected for a pilot tumor study, signaling sufficient backcrossing. For macrophage-targeted Eya3 knockout, we crossed Eya3-flox-C57BL/6 mice with MafB-Cre mice[56] (The Jackson Laboratory RRID:IMSR_JAX:029664), which provides additional C57BL/6 DNA. The following primer sequences were used to genotype mice:

Eya3-flox Forward: CCA CTT GGA GTA AGC ATC CAG TC
Eya3-flox Reverse: AAG CAA AAC GTC CTA GGT GCT C

The WT and flox-Eya3 allele can both be amplified by this pair of primers.

MafB Common Forward: AGA ACG AGA AGA CGC AGC TC
MafB Reverse: GGC GCA GAA TAG GGA GTC T
MafB-Cre Reverse: GCG CAT GAA CTC CTT GAT GA

The MafB WT allele is amplified through a PCR reaction with “MafB Common Forward” and “MafB Reverse”. The MafB-Cre allele is amplified through a PCR reaction with “MafB Common Forward” and “MafB-Cre Reverse”.

All genotyping PCR reactions were amplified using GoTaq Green (Promega #M7122, Madison, WI) and run on 3% agarose gels. Genotypes utilized throughout this paper include Eya3fl/fl (flox control), MafB-Cre (Cre control), and Eya3fl/fl;MafB-Cre (macrophage Eya3 homozygous knockout).

### RNA-seq analysis

RNA was extracted from macrophage samples using the RNeasy RNA Isolation Kit (Qiagen) in triplicate, then sequenced by the University of Colorado Cancer Center Genomics Core Facility. Libraries were prepared using the NuGEN Universal Plus mRNA-seq kit, and samples were sequenced with paired-end reads (40 million reads/sample) on an Illumina NovaSEQ6000 sequencing instrument. RNA-seq data processing was performed utilizing the nf-core/rnaseq pipeline (v. 3.12.0)[77]. Raw read quality was evaluated utilizing FastQC and MultiQC as part of the pipeline’s standard quality control procedures. Illumina adapters and low-quality reads were subsequently removed using Trim Galore! (v. 0.6.7). Trimmed sequence reads were aligned to the murine reference genome (GRCm39, Ensembl release 104) utilizing STAR (v. 2.7.4)[78]. Transcript-level quantification was performed utilizing Salmon[79] against an index incorporating decoy sequences and non-coding RNAs. Differential gene expression analysis was performed utilizing the DESeq2 R package (v. 1.48.1)[80] Pathway enrichment analysis was performed using GSEA[81]with the following collections of gene sets: Hallmark, Reactome, WikiPathways, and Gene Ontology (GO) Biological Processes. Heatmaps were generated using the ComplexHeatmap R package (2.24.1)[82]. They display representative genes associated with the indicated biological processes and were selected to illustrate transcriptional changes within pathways identified by GSEA. These genes were not restricted to GSEA leading-edge genes. Raw data can be found in the Gene Expression Omnibus database under accession number xxxxx (data to be deposited soon).

### Generation of bone marrow derived macrophages (BMDMs)

Bone marrow was harvested from the femurs and tibias of humanely euthanized mice through centrifugation. The resulting bone marrow pellet was washed in macrophage media, passed through a 70uM filter, then plated in macrophage media with 0.02ng/uL macrophage colony stimulating factor (M-CSF) (Sigma cat#M9170-10ug). Media and M-CSF were refreshed every 2-3 days, and BMDMs were used for their intended purpose within approximately a week from the time bulk bone marrow was plated. BMDMs were grown at 37°C in 5% CO_2_.

### Antigen processing assay

Macrophages were plated in a 6-well format, and the assay was performed when cells reached confluence. DQ-Ovalbumin (DQ-OVA) antigen processing reagent (ThermoFisher Scientific #D12053, Waltham, MA) was used at a concentration of 5ug/mL for the desired time (1.5hrs for RAW246.7 cells and BMDMs, and 30min for J774A.1 cells) at 37°C under normal incubator conditions. A negative control well (no DQ-OVA), and an ice control well (DQ-OVA added to cells placed on ice) were used for each experiment. Cells were fixed with 10% buffered formalin for 10min at endpoint, washed, resuspended in FACS buffer (1% FBS in Hanks’ Balances Salt Solution (HBSS)), or not fixed, and stained with propidium iodide as a live/dead marker in antigen processing assays utilizing 72hr, 2uM LG1-34 treatment. Samples were analyzed on a BD Accuri flow cytometer using Flowjo software (TreeStar, Ashland, OR).

### Antigen presentation assays

CD8 (OT-1) T cells were isolated using the MojoSort CD8+ T cell isolation kit (Biolegend) and labeled with violet proliferation dye (VPD) (BD Biosciences), per manufacturer’s instructions, prior to co-culture. M-CSF-matured BMDMs were isolated and cultured as described and were treated with LPS (200ng/mL) and either ovalbumin or SIINFEKL (5 ug/mL) for 12 hrs at 37°C. Following treatment, cells were washed and then co-cultured with isolated OT-1s for three days at a 1:10 ratio of BMDM:OT1. Cells were then washed, stained and run on a flow cytometer.

### Flow cytometry analysis of antigen presentation assays

Cells were stained for surface markers with anti-mouse CD8 antibody (Biolegend, clone 53-6.7) PD-1 (Biolegend, clone 29F.1A12) CD44 (Tonbo, clone IM7) CD45 (Biolegend, clone 30F-11), B220 (Biolegend, clone RA3-6B2) for 30 minutes at 37°C. Cells were washed and fixed with 1% paraformaldehyde and 3% sucrose for 10 mins in the dark at room temperature. Cells were washed twice with FACS buffer (0.1% BSA, 1x HBSS, 2mM ethylene diamine tetra acetic acid and 0.02% sodium azide). Cells were resuspended in FACS buffer before acquiring on Cytoflex LX. Analysis of responding cells to assay T cell proliferation index was performed with Flowjo software and calculated with the following equation:

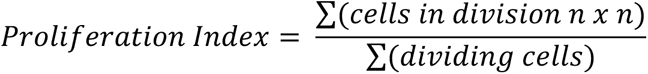

### Cloning, expression, and purification of Eya3-ED

The mouse Eya3-ED (residues 225–573) was cloned into GEX-6P-1 (GE Healthcare) with an N-terminal GST tag and a PreScission protease cleavage site. Constructs were transformed into *E. coli* BL21(DE3) cells. Cells were grown at 37°C until OD_600_ reached 0.6–0.8. Protein expression was induced with 0.3–0.4 mM IPTG at 18–20 °C for 16–20 h. Cells were harvested by centrifugation. Cell pellets were resuspended in lysis buffer [50 mM HEPES pH 7.5, 300 mM NaCl, 5% (v/v) glycerol, 1 mM DTT, 0.2 mg/mL lysozyme (ReadyLyse, Lucigen), 25 U/mL Benzonase (Sigma), and 1mL of 100x Xpert Protease Inhibitor Cocktail Solution (GenDEPOT) per 100mL buffer. Resuspended cells were lysed by sonication. After adjusting NaCl concentration to 500mM, the lysates were clarified by centrifugation and then filtered using a 0.45 µm filter. Clarified lysates were incubated with glutathione sepharose beads (Cytiva) for 1 hr at 4°C, with flowthrough reapplied once to maximize binding. Beads were washed with 25 CV of lysis buffer, then incubated overnight at 4°C with PreScission protease to cleave the GST tag. Cleaved protein was eluted with buffer [50 mM HEPES pH 7.5, 300 mM NaCl, 5% glycerol, 1 mM DTT], concentrated and applied to a Superdex 200 Increase 10/300 GL (Cytiva). Fractions corresponding to monomeric Eya3-ED were pooled, concentrated to 10-15mg/mL, and flash frozen in aliquots stored at -80 °C.

### Thermal stability assay (TSA)

Thermal stability assays were performed to evaluate compound binding dependent stabilization of purified Eya proteins. Prior to the assay, recombinant proteins stored in glycerol-containing buffers were buffer exchanged into glycerol-free TSA assay buffer (containing 50 mM Tris-HCl pH 7.5, 250 mM NaCl, 1.5 mM TCEP, and 500 µM MgCl_2_). Buffer was stored at 4°C and used within one month. Eya-PTP phosphatase inhibitor compound LG1-34 was prepared by diluting 10 mM stocks to 2 mM in 100% DMSO. DMSO alone was used as the vehicle control. Following buffer exchange, recombinant Eya3-ED was normalized to 0.5 mg/mL in glycerol-free TSA assay buffer.

For each treatment group, 0.4 µL of 2 mM compound stock or DMSO was added to a PCR tube. Normalized recombinant Eya3-ED protein was then added to a final volume of 40 µL, yielding a final compound concentration of 20 µM. After compound incubation on ice, 20 µL of each reaction was transferred to a labeled low-retention microcentrifuge tube and kept on ice as the unheated control. The remaining 20 µL was heated in a thermocycler for 3 min at the optimized assay temperature. A temperature of 50°C was used for single-temperature screening, while a range of 38.5–62.5°C was used when determining apparent melting behavior. Following heating, samples were allowed to equilibrate at room temperature for 3–5 min to promote uniform aggregation and prevent refolding of thermally denatured proteins upon placing on ice. Samples were then transferred to low-retention microcentrifuge tubes and centrifuged at 15,000 rpm for 1hr at 4°C. 10 µL of the soluble supernatant was carefully collected without disturbing the pellet, and GAPDH was spiked in as a loading control. Samples were analyzed on SDS-PAGE gels which were stained with SYPRO Ruby total protein staining solution and visualized using a Bio-Rad ChemiDoc MP Imager.

### Animals and *in vivo* tumor studies

Animal studies were performed in accordance with protocols approved by the Institutional Animal Care and Use Committee (IACUC) at the University of Colorado Anschutz Medical Campus. Only female mice were used for breast cancer studies, as breast cancer primarily affects females (over 99% of cases). Male and female mice were used for *ex vivo* studies with BMDMs to maximize use of bred animals.

For tumor studies in C57BL/6J in Eya3fl/fl, Eya3fl/fl;MafB-Cre, and MafB-Cre mice (bred in house), 500,000 E0771 cells in serum-free media were injected into the fourth mammary fat pad of 11 to 13-week-old mice, age-matched between control and knockout groups. Tumors were measured by caliper 3 times a week. Mice were euthanized at an ethical endpoint of 1.5cm^3^ for generation of a Kaplan-Meier survival curve (study #1), or all at once on day 18 (study #2)

For tumor studies with small molecule Eya tyrosine phosphatase inhibitor LG1-34, 1x10^6^ E0771-gfp-luc cells in serum-free media were injected into the fourth mammary fat pad of 8-week-old, immune competent C57BL/6J mice (The Jackson Laboratory RRID:IMSR_JAX:000664) or 500,000 E0771-gfp-luc cells were injected into the fourth mammary pad of 8-week-old, immune compromised NSG mice (The Jackson Laboratory RRID:IMSR_JAX:005557). Mice were imaged and weighed at the noted intervals (every 3 days for C57BL/6 study #1, twice a week for C57BL/6 study #2, twice a week for NSG immune compromised study). Mice were treated daily by oral gavage with 50mg/kg LG1-34 in carboxymethylcellulose (CMC), or an equivalent volume of DMSO in CMC. Imaging was performed under isoflurane anesthesia on the IVIS200 bioluminescent imager 10 min after intraperitoneal injection with 100uL of 100x luciferin (Gold Biotechnology, LUCK-1G, St. Louis, MO), and images were analyzed with Living Image software. Luminescence is reported as total flux (p/s) of the tumor minus total flux (p/s) of background. Mice were euthanized all at once on day 23 (C57BL/6 study #1), once tumors reached 1cm^3^ (C57BL/6 study #2), or all at once on day 19 (NSG study).

### Multiplexed IHC of fixed tumor samples

Mouse tumors from *in vivo* experiments were fixed in 10% buffered formalin and formalin-fixed paraffin-embedded (FFPE) blocks were generated. N=4 size matched tumors per group were fixed, and two slices per tumor were stained from C57BL/6 macrophage Eya3 KO tumor study #2, and N=6 tumors per group (one slice per tumor) were stained from LG1-34 tumor study #1. Tumor slices from FFPE blocks were stain with a macrophage and T cell focused antibody panel (F480, CD80, CD163, CD3, CD8, EGFR, DAPI) by the Human Immune Monitoring Shared Resource (HIMSR) at the University of Colorado Anschutz Medical Campus, and imaged on the Akoya Biosciences’ Vectra Polaris, then segmented and preliminarily analyzed using inForm image analysis software by HIMSR. Final data were viewed with the Enable-Medicine Visualizer tool.

### Advanced spatial analysis of tumor slices

Multiplexed tumor images were passed through an advanced spatial analysis pipeline, in accordance with the techniques described in depth by Bouchard et al. [60]. In brief, cell populations were assigned with the CELESTA machine learning algorithm (https://github.com/plevritis-lab/CELESTA), which can be informed by a matrix of anticipated cell populations based on marker combinations, but can also identify populations with unexpected marker combinations. All CELESTA assignments were manually verified by visualizing cell assignments in comparison to the original immunofluorescent images (Supp. Fig. 11A, 14A), and adjusting marker thresholds as appropriate. The matrix of marker identities of cell populations in our analysis can be found in Supp. Table 1. Expression thresholds are found in Supp. Tables 2-3.

To quantify immune cell spatial organization within tumors, we performed colocatome analysis that utilizes the colocation quotient (CLQ) spatial metric[59, 60]. CLQ can be used to quantify how a specific cell population spatially co-locates with another cell population within its nearest neighbors (set as 20), and was calculated between all cell populations identified, in a pairwise manner, using the equation: CLQ_b→a_ = (C_b→a_/N_a_) / (N_b_/(N − 1)) where C_b→a_ is the number of cells of cell type b among the defined nearest neighbors of cell type a, N is the total number of cells and N_a_ and N_b_ are the numbers of cells for cell type a and cell type b, respectively. Significance of CLQs within each tumor image is assessed using spatial permutation analysis to determine non-random cell population pair organization. Supplemental Excel Files 1 and 2 contain all supplementary CLQ data.

### Analysis for macrophage and CD8+ T cell localization within tumors

CELESTA images of macrophages and CD8+ T cells were analyzed using ImageJ software. The “peripheral” region of the tumor was defined as the region that spans from the outer tumor boundary to a region that is just within the tumor boundary, that is 10% of the average diameter of the tumor. The “central” region of the tumor is defined as everything internal to the “peripheral” region. Macrophages and CD8+ T cells were masked within these regions, and the ratio of the area of masked cells (central vs. peripheral) was recorded.

### Flow cytometry

To prepare blood for flow cytometry analysis, 100-200uL of blood was collected in 500uL of 2uM ethylenediaminetetraacetic acid (EDTA)-HBSS through submandibular bleed, followed by red blood cell lysis with 1x Ammonium-Chloride-Potassium (ACK) buffer for 5min prior to staining. Mice were then humanely euthanized, tissues (tumor, lung, mammary gland, lymph node, spleen) were promptly removed and placed into immune cell media (RPMI + 10% FBS + 1% Pen/Strep +50 μM β-mercaptoethanol) on ice. Tumors, lungs, and mammary glands were minced and incubated in digestion buffer (1× HBSS, collagenase IA (0.1 mg/ml), and DNase I (60 U/ml) at 37°C for 25 min with continuous shaking. All tissues were then mechanically dissociated by passage through a 70uM nylon filter, then immediately stained for flow cytometry, or frozen in complete medium with 10% DMSO at -80°C for later staining.

Single cell suspensions were incubated with Zombie Fixable Viability Dye (BioLegend #423101, San Diego, CA) for 10 min, then washed and stained for 30-45min with fluorophore-conjugated antibodies (Supp. Table 3). Cells were fixed using the BD Cytofix/Cytoperm Fixation/Permeabilization Kit (BD Biosciences #554714, Franklin Lakes, NJ), and analyzed on a Bio-Rad ZE5 Cell Analyzer through the University of Colorado Cancer Center Flow Cytometry Shared Resource.

### Analysis of the association of macrophage Eya expression with clinical outcomes in TNBC patients treated with immune checkpoint inhibitors

#### Kaplan-Meier plots (GSE241876)

Bulk RNA-seq and clinical data from a single-arm phase II trial of carboplatin, nab-paclitaxel, and pembrolizumab in metastatic TNBC[66] were retrieved from GEO (accession GSE241876). Analyses were restricted to one baseline (pre-treatment) biopsy per patient (n=15). Expression values from the deposited DESeq2-normalized count matrix for *EYA1–4* were log2(normalized count + 1) transformed. For each gene, patients were dichotomized at the median expression value, and overall survival (OS) and progression-free survival (PFS) were compared between high- and low-expression groups by the Kaplan-Meier method with two-sided log-rank tests. Composite *EYA* scores were computed as the mean of per-gene z-scores (*EYA1*+*EYA3*) across the 15 baseline samples and evaluated using the same median-split procedure. All analyses were exploratory and unadjusted for multiple comparisons. Survival analyses and figures were generated in R using the survival and ggplot2 packages.

#### UMAP (GSE246613)

Processed scRNA-seq objects from a phase Ib/II trial of neoadjuvant pembrolizumab followed by pembrolizumab plus stereotactic radiation therapy in early-stage TNBC[65] were obtained from GEO (accession GSE246613). The dataset comprised separately deposited CD45+ immune (342,749 cells) and non-immune (171,746 cells) compartments. Precomputed UMAP coordinates and cell-type annotations provided by the authors were used without re-clustering or re-embedding, with cell identities displayed at a coarse resolution (immune: T cells, B cells including plasma cells, myeloid cells, and mast cells; non-immune: cancer cells, normal epithelium, iCAFs, myCAFs, endothelial cells, and perivascular-like cells). Expression values were normalized to 10,000 counts per cell and log1p-transformed (natural log), and EYA1–4 expression was overlaid onto the UMAP embeddings, with cells plotted in ascending order of expression to prevent masking of positive cells. Color scales are gene-specific and not directly comparable across genes. Visualizations were generated in R using ggplot2.

#### Supplemental detection of Eya frequency in immune cells and association between Eya expression and tumor grade

The integrated breast tumor scRNA-seq atlas from Xu et al.[50]; comprising 236,363 treatment-naive cells across eight public studies, was obtained from the authors’ data deposit (Zenodo, DOI: 10.5281/zenodo.10672250) with the authors’ harmonized cell-type annotations. For each EYA family gene (EYA1-4) and major cell lineage, detection frequency was defined as the percentage of cells with non-zero expression (>= 1 UMI count or >0 log-normalized value, which are mathematically equivalent). Detection bar plots were generated in Python using matplotlib and seaborn. For tumor grade associations, macrophages from patients with documented histological grade were aggregated to patient-level mean expression (mean log-normalized counts per patient; patients with <10 graded macrophages were excluded, leaving n=57 patients). Associations between EYA expression and numeric tumor grade were evaluated by two-sided Spearman rank correlation. Box plots with overlaid patient-level data points were generated in R using ggplot2.

#### Macrophage Eya association with clinical-response (GSE169246)

scRNA-seq profiles of CD45+ immune cells from paired tumor and blood specimens of 22 patients with advanced TNBC treated with paclitaxel +/- atezolizumab[67] (GEO accession GSE169246) were re-analyzed from deposited raw count matrices. Data processing including quality control filtering, library size normalization, Leiden graph-based clustering, and marker-based lineage annotation was performed in Python using scanpy. The myeloid compartment was subsequently resolved into macrophages and monocytes by marker-based label reassignment using an expanded marker panel, without re-clustering. For each patient, raw UMI counts from tumor-infiltrating macrophages (or monocytes) were summed across the specified timepoints (either all timepoints pooled, or pre- and on-treatment samples with progression biopsies excluded) and normalized to counts per 10,000 (CP10k) followed by log1p transformation (natural log) to yield patient-level pseudobulk expression. Pseudobulk EYA1–4 levels were compared between clinical responders (partial response [PR]) and non-responders (stable disease [SD] or progressive disease [PD]), as recorded in the trial’s supplementary clinical table, using two-sided Mann-Whitney U tests. Subsets containing fewer than two evaluable patients or lacking detectable expression were categorized as untestable. Figures were generated in Python using matplotlib.

### Statistics

Prism software (v10.0; GraphPad) was used for all statistical analyses. Two-tailed unpaired Welch’s t-test was used when comparing two groups. One-way Analysis of Variance (ANOVA) was used when comparing 3 groups, without the element of time. Two-way ANOVA was used to analyze the significance of tumor growth over time (p values reported are “Time x Column Factor”). Spatial permutation (n = 500) was used to assess significance of CLQs. All *in vitro* experiments were run with at least 3 technical replicates and repeated with at least 3 biological replicates.

## Acknowledgments

We thank the Breeder Core at AMC for their assistance breeding mice for the generation of the macrophage Eya3 conditional knockout model. We thank the Human Immune Monitoring Shared Resource (RRID:SCR_021985) within the University of Colorado Human Immunology and Immunotherapy Initiative and the University of Colorado Cancer Center (P30CA046934) for expert assistance with fluorescent IHC multiplex imaging. This work also used the Cell Technologies (RRID:SCR_021982), Genomics (RRID:SCR_021984), Biostatistics and Bioinformatics (RRID:SCR_021982), and Flow Cytometry (RRID:SCR_022035) shared resources at the University of Colorado Anschutz Medical Campus and the University of Colorado Cancer Center (P30CA046934). Special thanks to C. Jakubzick, K. Schwertfeger, and L. Fry for their expertise on macrophage biology and specialized assays, and to Tal Keidar for her pathology assessment of EGFR+ macrophages.

---

**Supplemental Figure 1.**
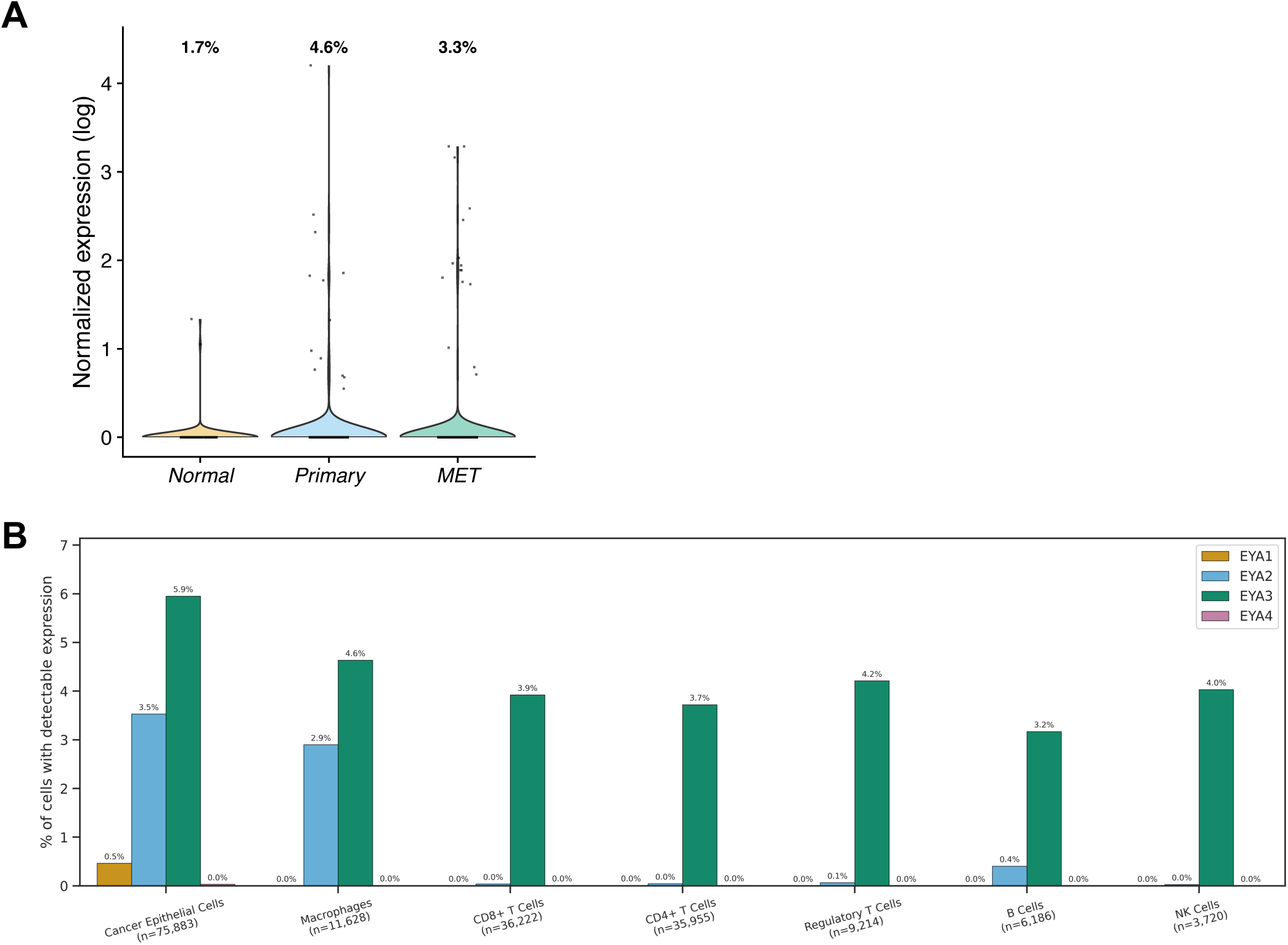
Additional plot for the quantification of Eya proteins in macrophages in murine normal breast, primary breast tumor, and lung metastasis, and Eya1-4 expression in TNBC and immune cells in an additional human TNBC study. A) Macrophage Eya3 expression in normal breast, primary tumor, and lung metastasis as a violin plot (GSE231915). B) Eya1-4 expression in TNBC cancer epithelial cells and immune cells across 8 public datasets[50].

**Supplemental Figure 2.**
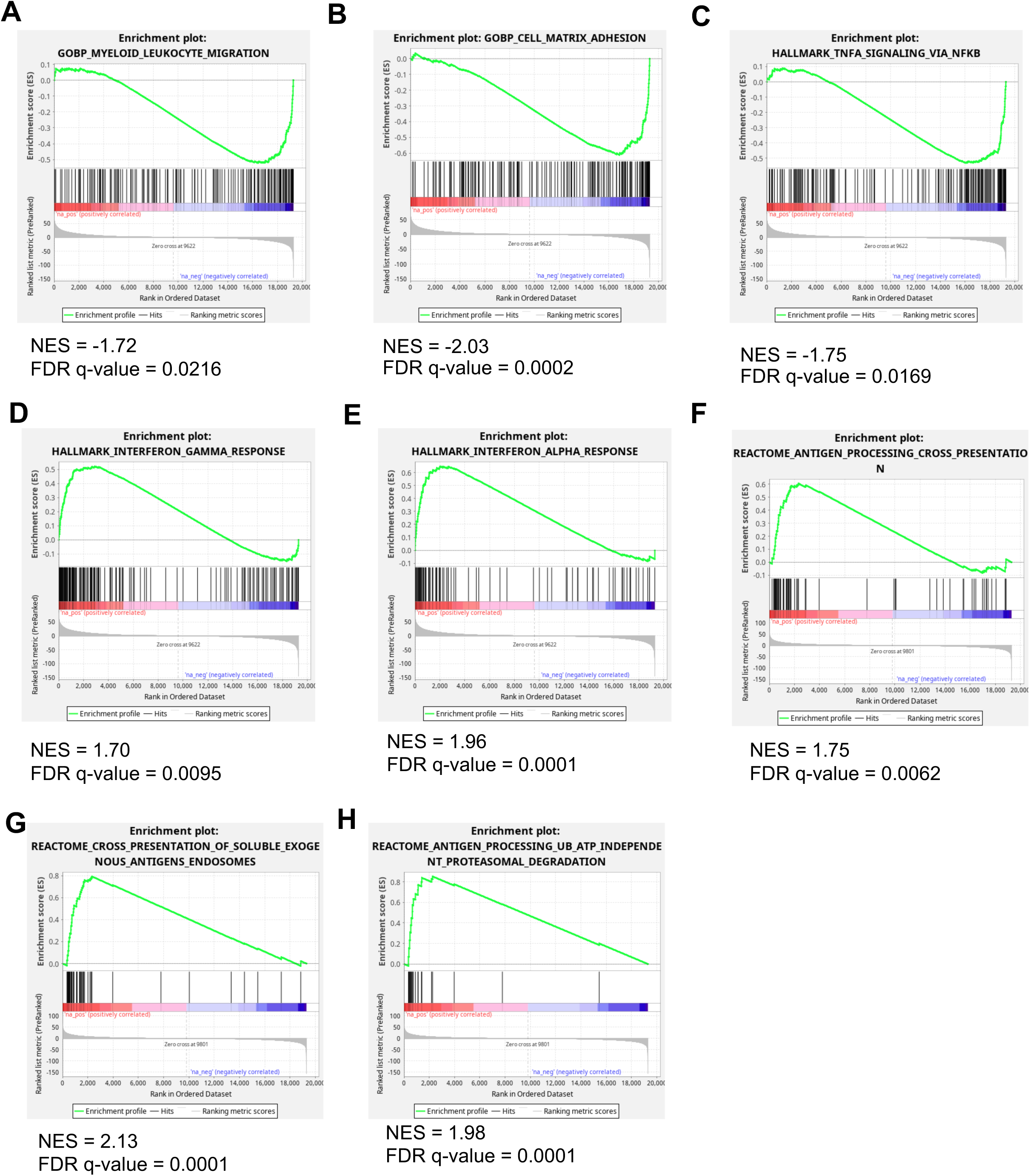
GSEA plots of pathways relevant to macrophage function from RAW264.7 shSCR and shEYA3 KD bulk RNA-seq. GSEA plots comparing RAW264.7 shEYA3 KD2 vs shSCR RNA expression: A) Myeloid Leukocyte Migration, B) Cell Matrix Adhesion, C) TNFA Signaling via NFKB, D) Interferon Gamma Response, E) Interferon Alpha Response, F) Antigen Processing and Presentation, G) Cross Presentation of Soluble Exogenous Antigens Endosomes, and H) Antigen Processing Ubiquitin ATP Independent Proteasomal Degradation.

**Supplemental Figure 3.**
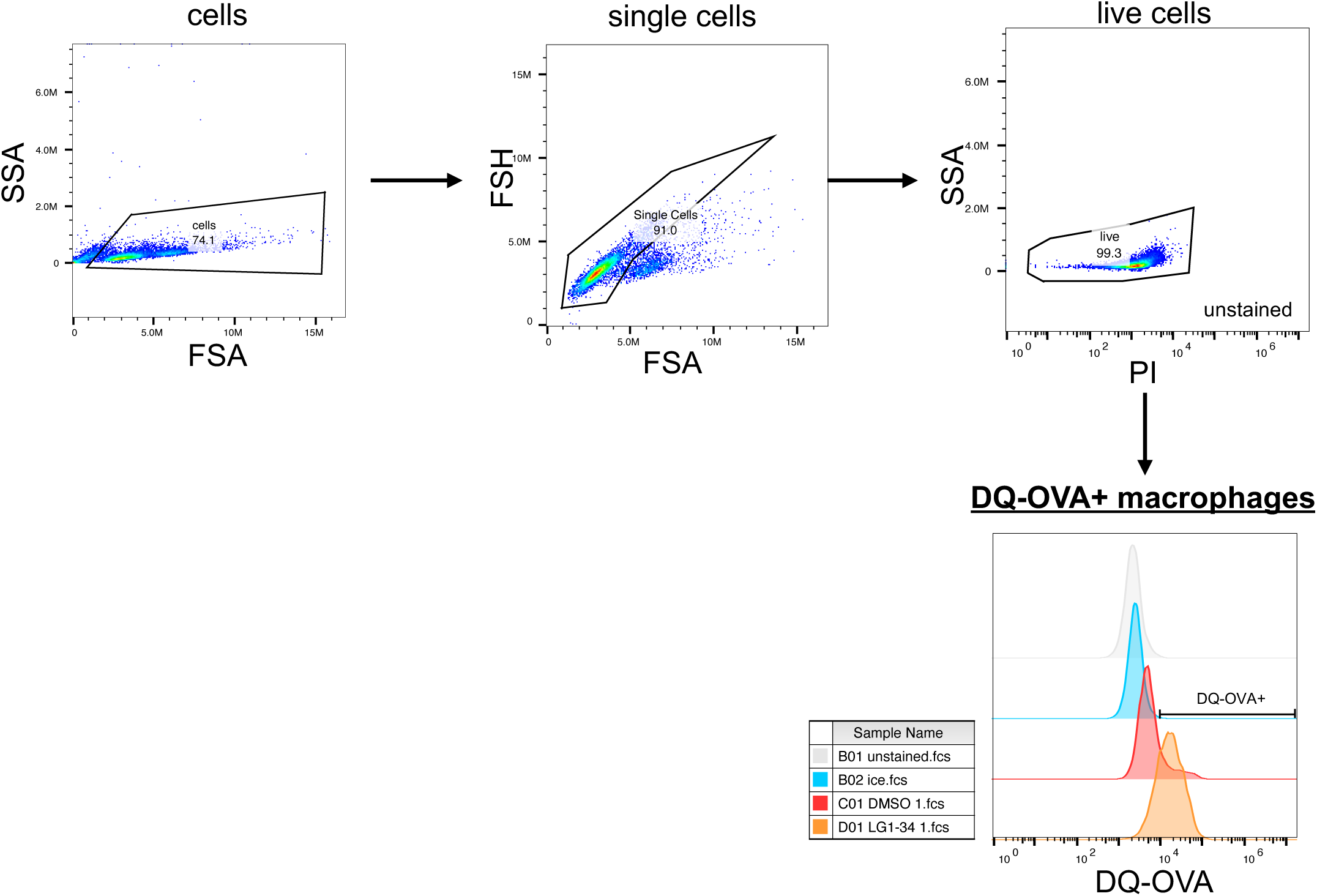
Gating strategy for DQ-OVA antigen processing experiments. Experiments represented in Fig. 2H-M, Fig. 3F-H, and Fig. 5H-M utilize this gating strategy.

**Supplemental Figure 4.**
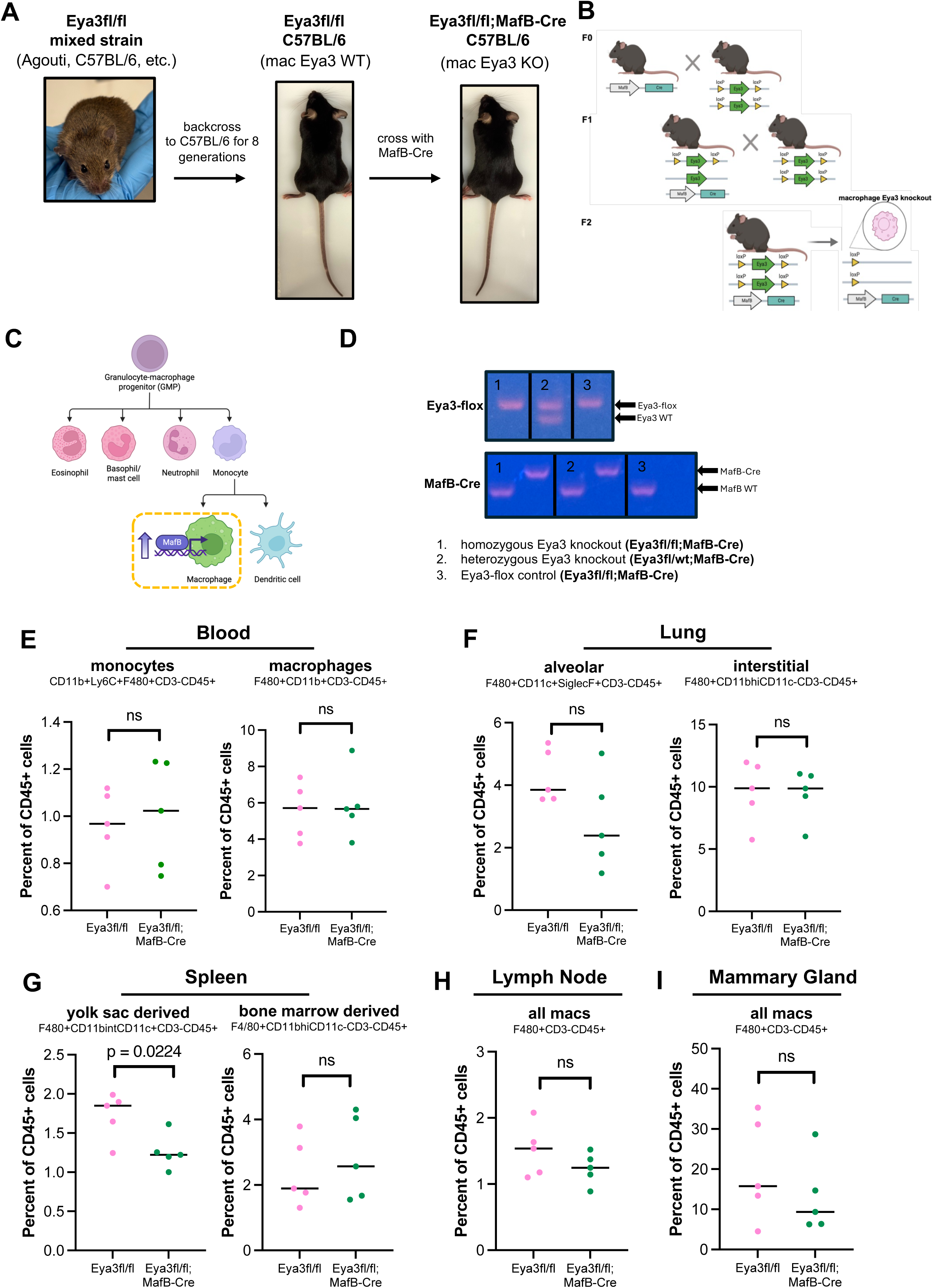
Additional characterization of the C57BL/6 macrophage Eya3 conditional knockout mouse. A) Strategy and images for developing a macrophage specific Eya3 KO mouse. Original Eya3fl/fl mixed strain mouse from R. Hegde lab. B) Complete breeding strategy for homozygous knockout of Eya3 in macrophages using MafB-Cre C57BL/6 and Eya3-flox C57BL/6 mice. C) MafB is upregulated in differentiated macrophages. D) Representative PCR products seen when genotyping littermates for Eya3 knockout status. E-I) Flow cytometric characterization of macrophage infiltration of tissues from control (Eya3fl/fl, n = 5) and Eya3 macrophage conditional knockout mice (Eya3fl/fl;MafB-Cre, n = 5). Statistical analysis performed using unpaired t-tests.

**Supplemental Figure 5.**
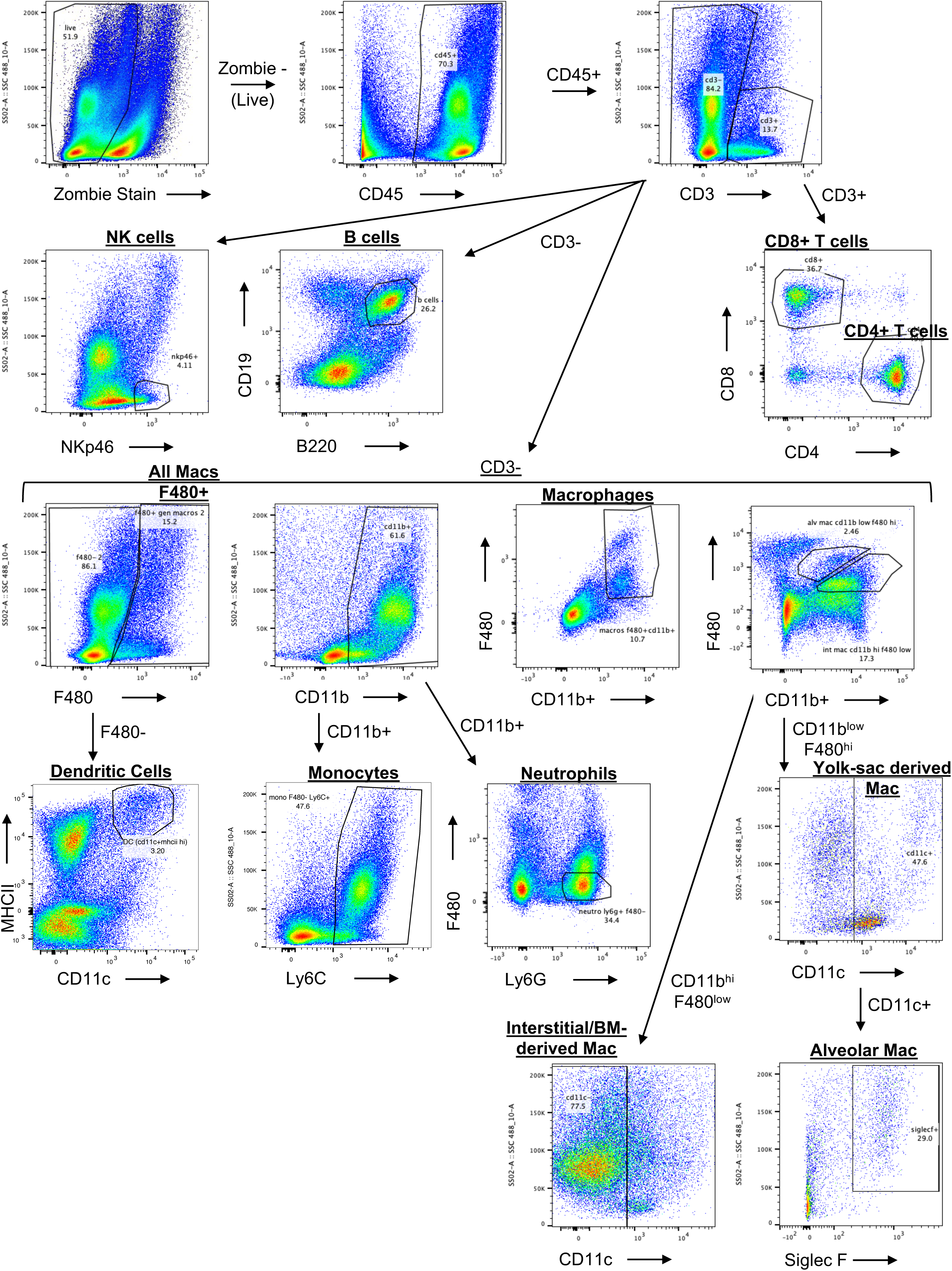
Gating strategy for identification of immune populations from organs and tumors of Eya3fl/fl and Eya3fl/fl;MafB-Cre mice. Gating strategy utilized to identify populations in Supp. Fig. 4E-I, Supp. Fig. 6, and Supp. Fig. 10.

**Supplemental Figure 6.**
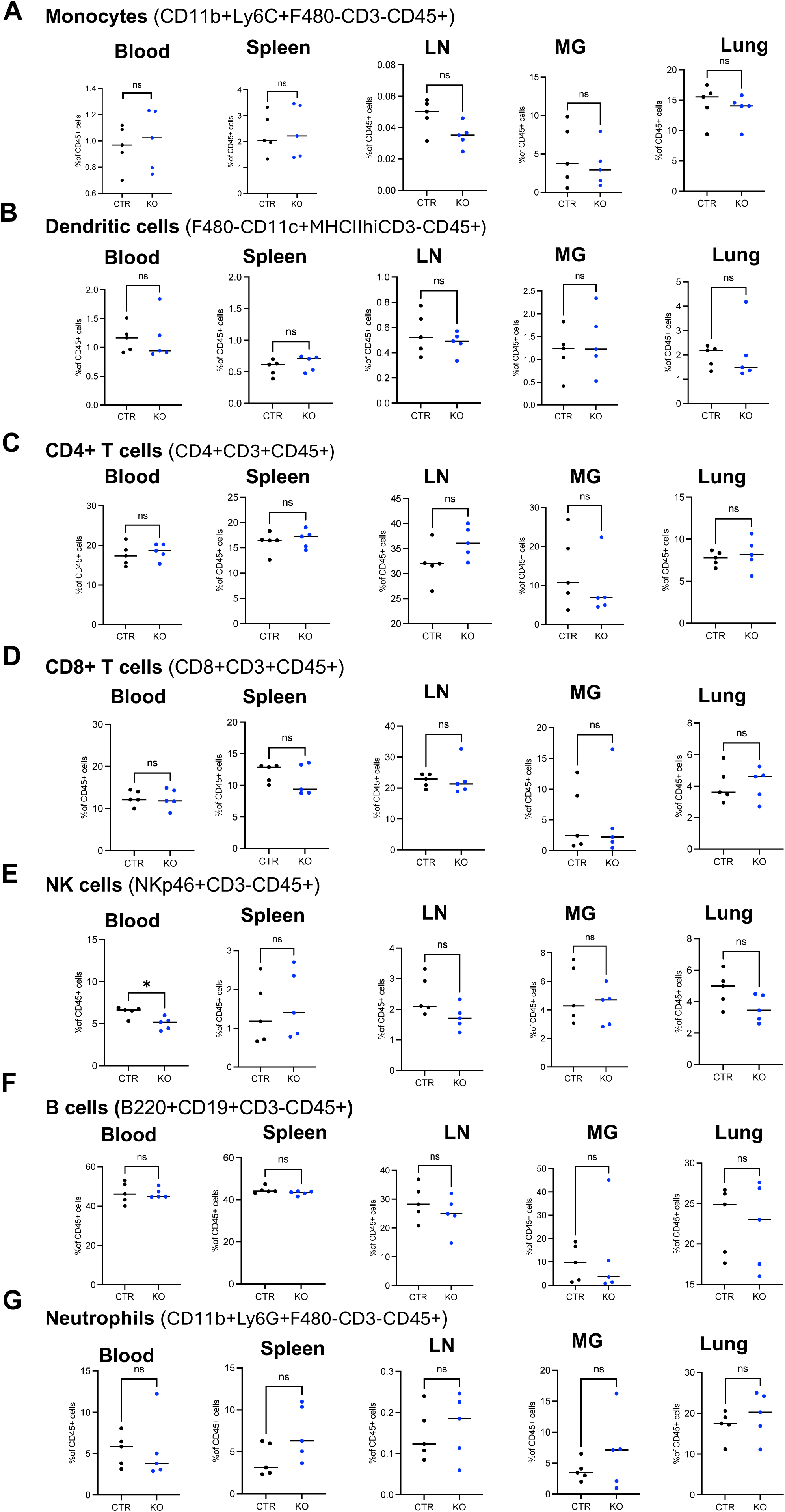
Quantification of additional immune cell populations within tissues in Eya3fl/fl (CTR) and Eya3fl/fl;MafB-Cre (KO) mice. Flow cytometry to examine: A) monocytes, B) Dendritic cells, C) CD4+ T cells, D) CD8+ T cells, E) NK cells, F) B cells, G) Neutrophils. Data from 5 Eya3fl/fl (CTR) and 5 Eya3fl/fl;MafB-Cre (KO) mice. Statistical analysis performed using unpaired t-tests. LN = lymph node, MG = mammary gland.

**Supplemental Figure 7.**
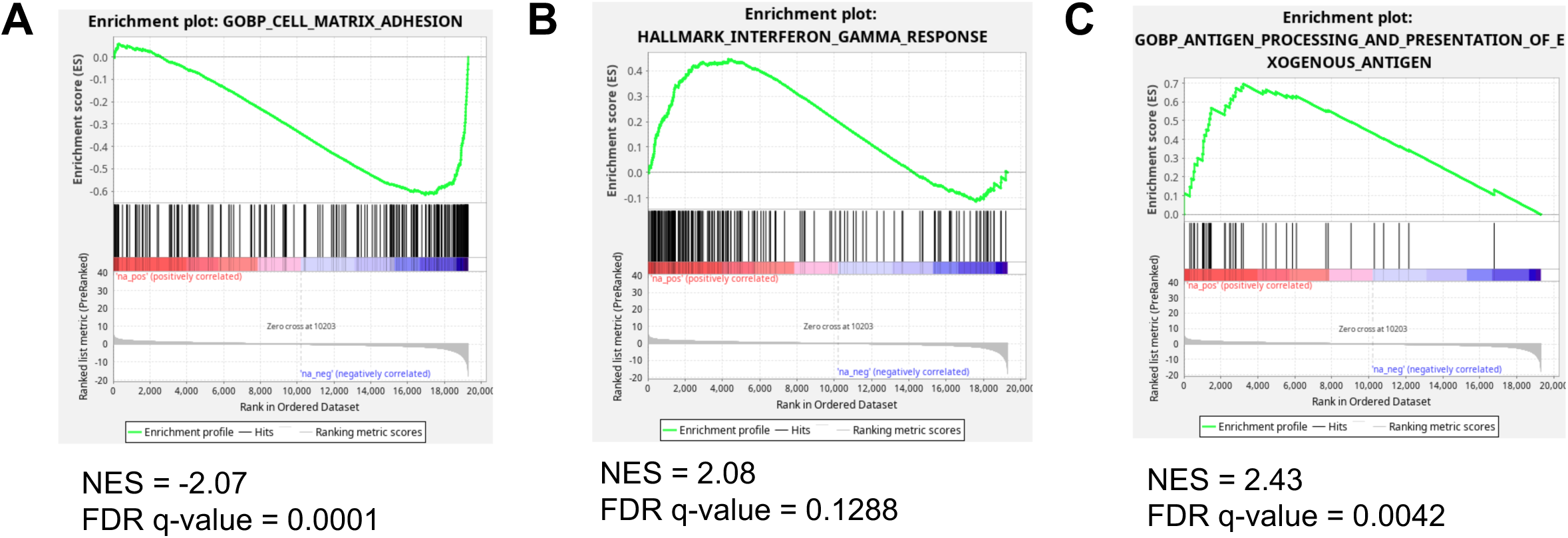
GSEA plots of pathways relevant to macrophage function obtained using RNA-seq of Eya3fl/fl (CTR) and Eya3fl/fl;MafB-Cre (KO) BMDMs. GSEA plots comparing RNA expression from BMDMs isolated from EYA3 KO vs CTR mice: A) Cell Matrix Adhesion, B) Interferon Gamma Response, E) Interferon Alpha Response, and C) Antigen Processing and Presentation of Exogenous Antigen. BMDMs were grown up from 1 mouse/per group in triplicate (3 independent wells).

**Supplemental Figure 8.**
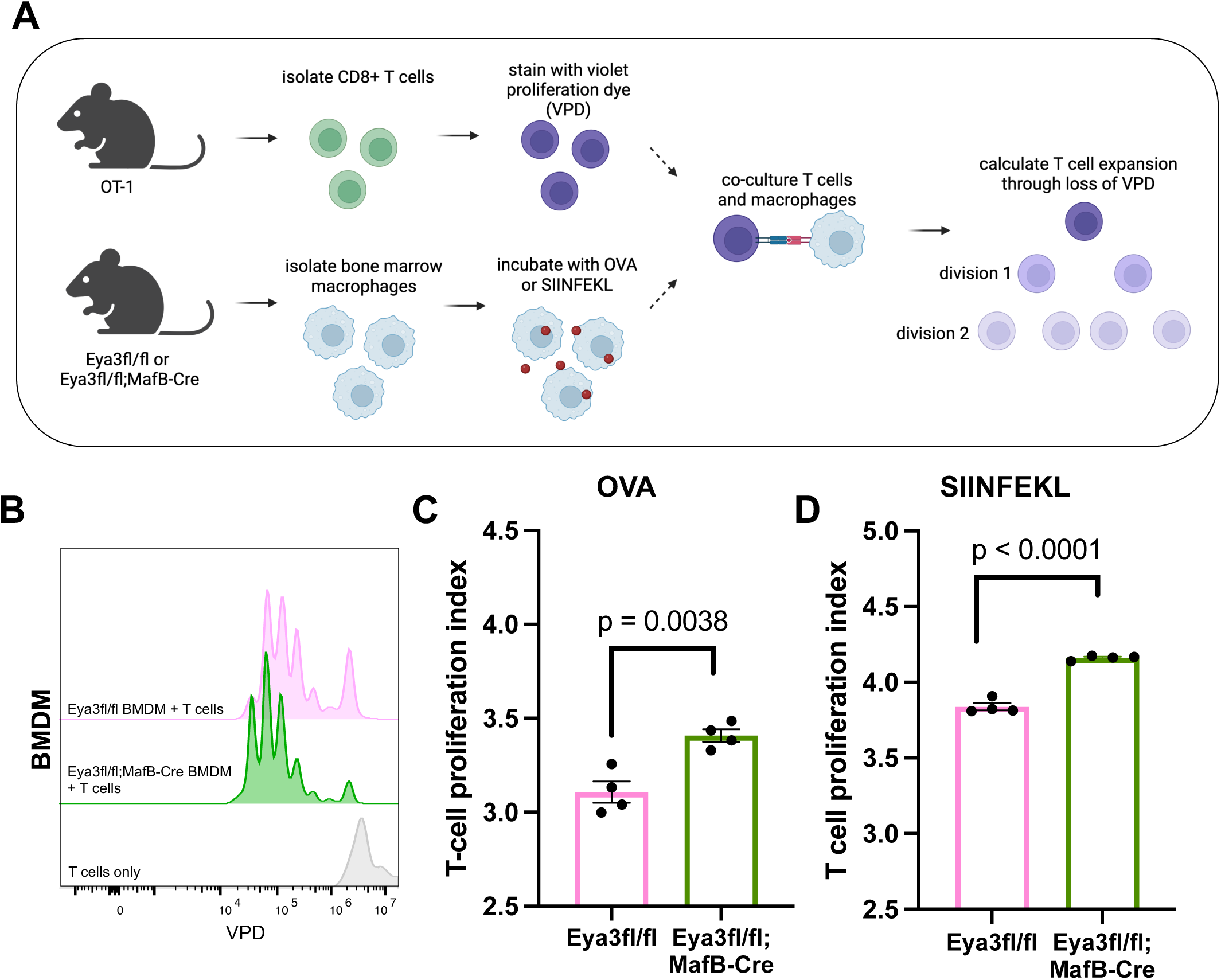
Loss of Eya3 modestly, but significantly, increases antigen presentation in BMDMs ex vivo. A) Schematic of antigen presentation assay. B) Representative histogram of flow cytometry from co-cultured CD8+CD44+B220-T cells used to evaluate VPD dilution. C) T cell proliferation index when BMDMs are primed with OVA. D) T cell proliferation index (calculated from VPD flow cytometry peaks) when BMDMs are primed with SIINFEKL. Statistical analysis performed using unpaired t-test. Experiments are representative of at least three biological replicates with technical triplicates.

**Supplemental Figure 9.**
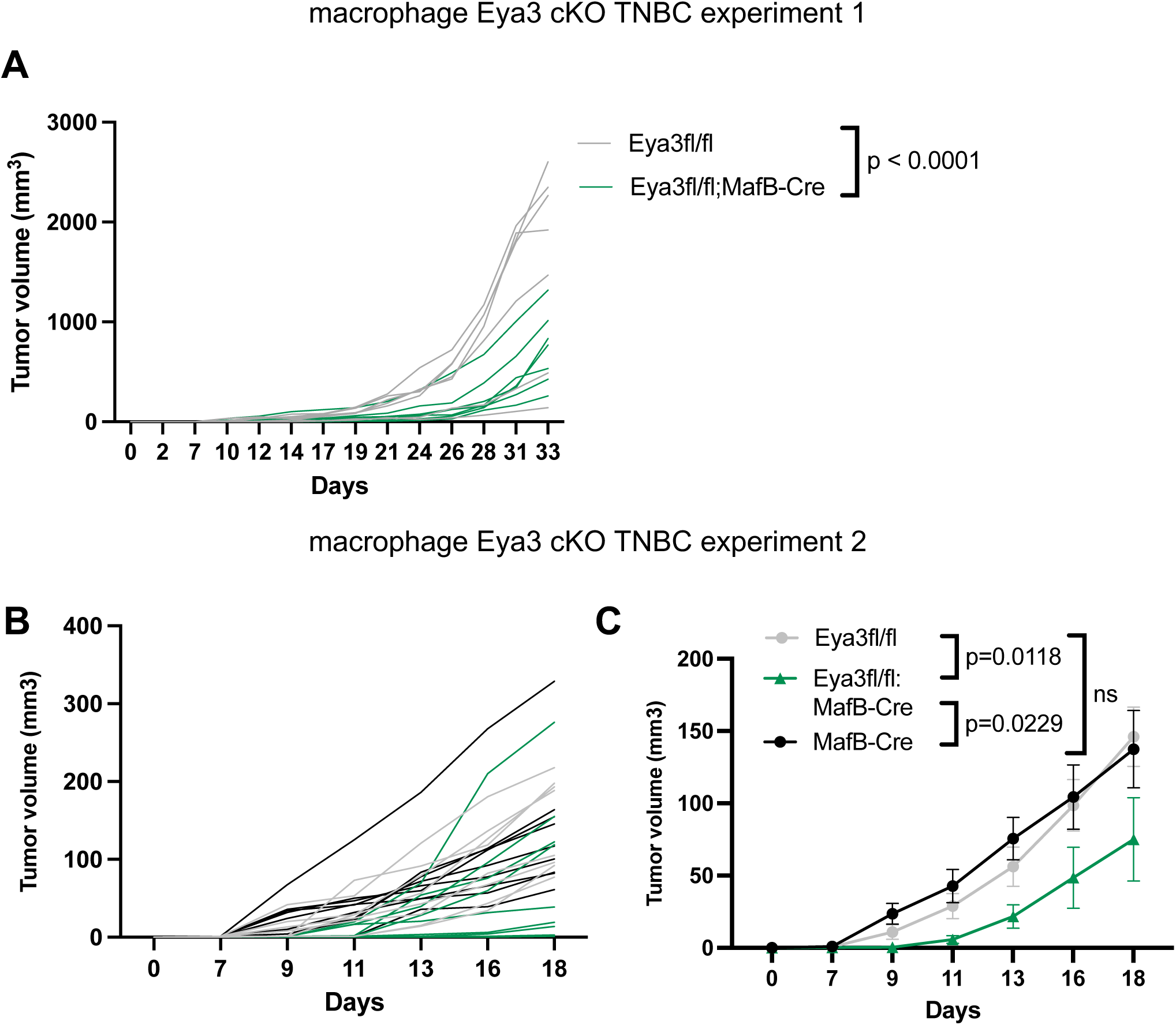
E0771 tumor growth in macrophage Eya3 conditional knockout mice. A) Macrophage Eya3 conditional knockout TNBC experiment #1: tumor growth tracks determined using caliper measurements for individual Eya3fl/fl mice (n = 7) and individual Eya3fl/fl;MafB-Cre mice (n = 7). B-C Macrophage Eya3 conditional KO TNBC experiment #2: B) Tumor growth tracks determined using caliper measurements for individual mice in experiment #2, C) Summary of growth data plotted for all 3 groups studied: Eya3fl/fl (Flox CTR, n = 8), MafB-Cre (Cre CTR, n = 9), Eya3fl/fl;MafB-Cre (Eya3 mac KO n = 10).

**Supplemental Figure 10.**
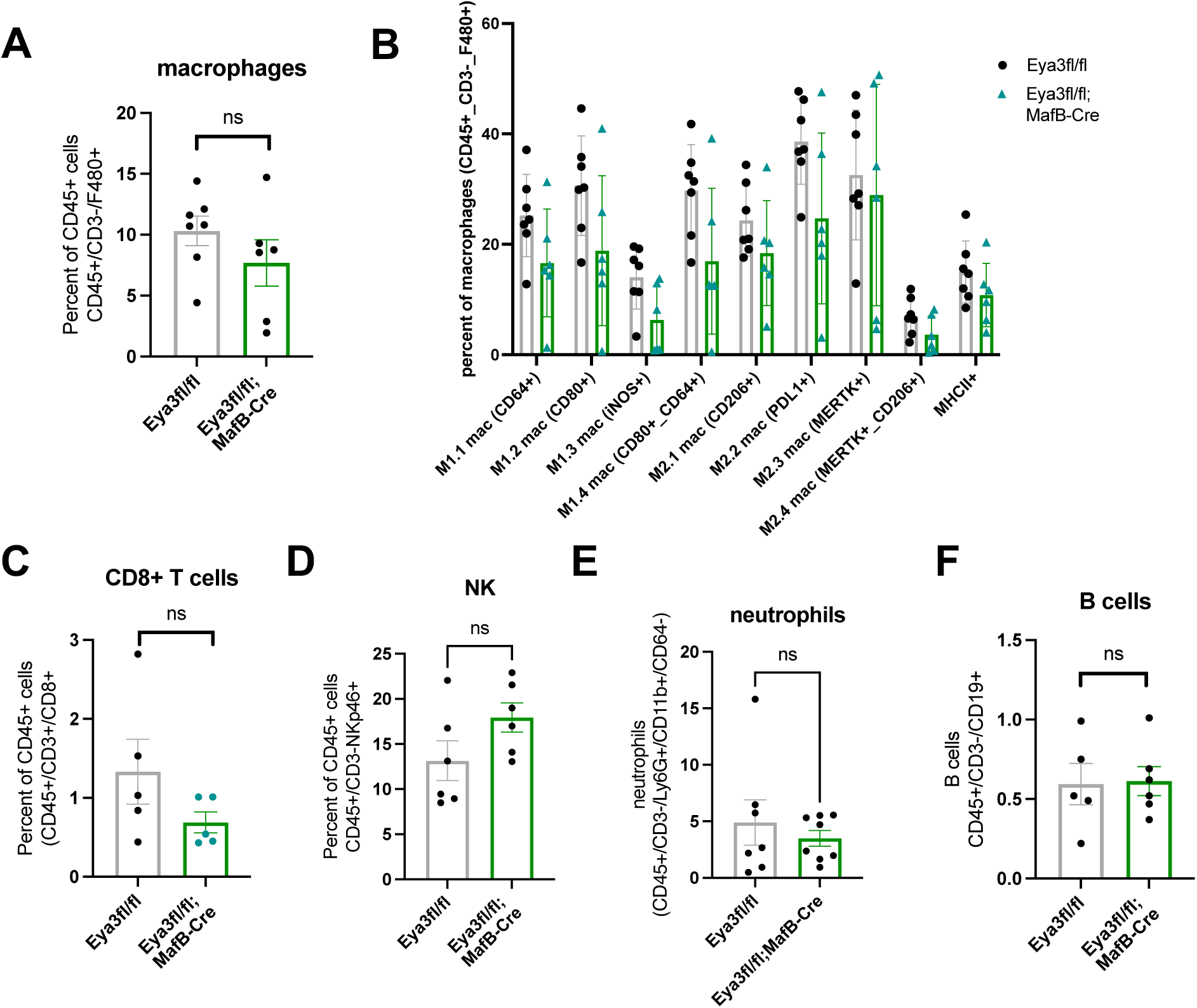
Flow cytometric analysis of immune infiltration of tumors from Eya3fl/fl (CTR) and Eya3fl/fl;MafB-Cre (KO) mice. Analysis performed on tumor experiment #1 to examine: A) total macrophages, B) Percent of macrophage subtypes, C) CD8+ T cells, D) NK cells, E) neutrophils, and F) B cells. Statistical analysis performed using unpaired t-tests.

**Supplemental Figure 11.**
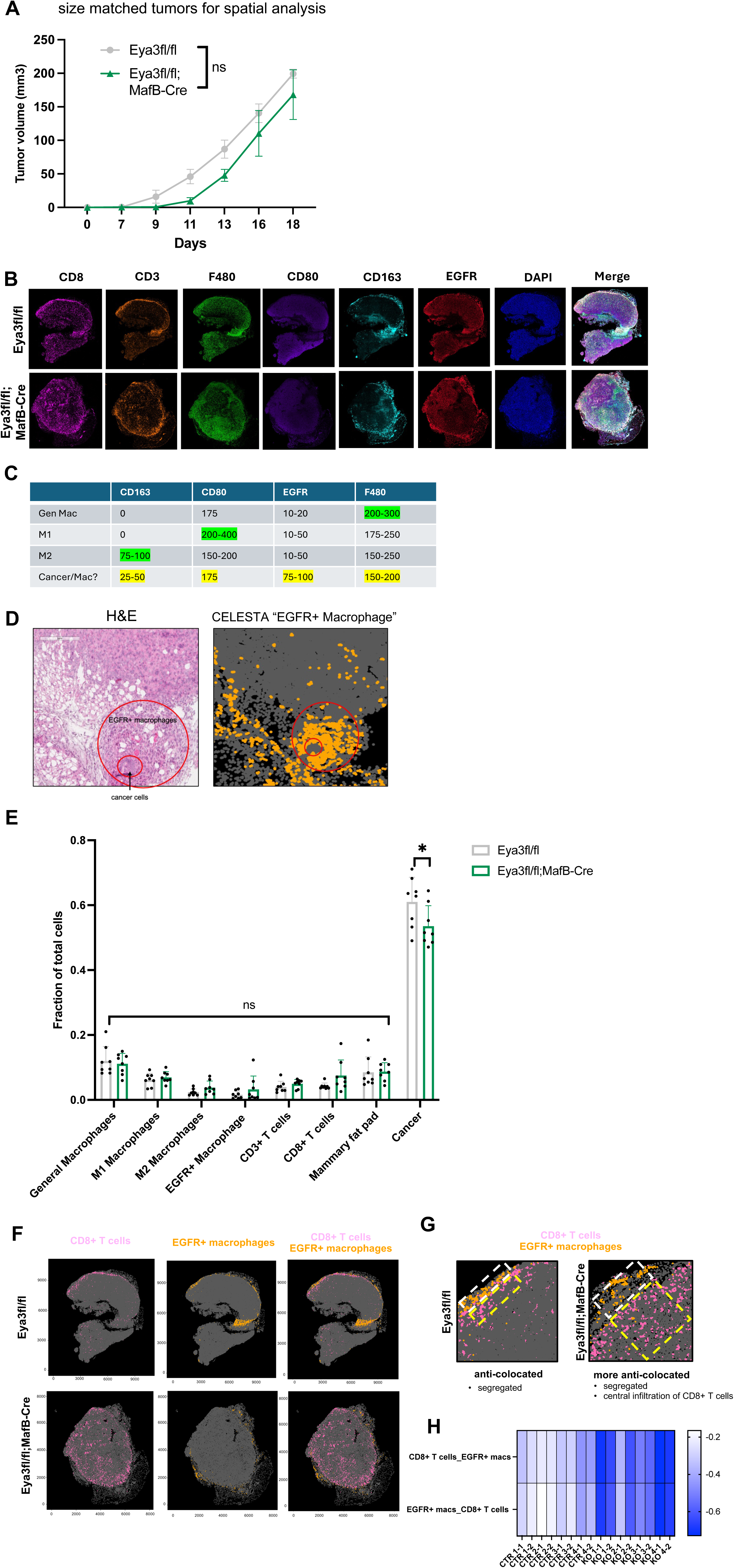
Supplementary material for spatial analysis of tumors from Eya3fl/fl (CTR) and Eya3fl/fl;MafB-Cre mice (KO). A) Summary tumor growth curves for size matched tumors (n = 4 for each group) analyzed in IHC spatial analysis. Statistical analysis performed using 2-way ANOVA. B) Fluorescent IHC images of all markers used to identify cell populations through CELESTA. C) Expression ranges of macrophage markers; unexpected macrophage population identified that expresses tumor marker “EGFR”. D) H&E staining demonstrating that the EGFR+ population is a macrophage subpopulation. Population was termed “EGFR+ macrophage”. E) Cell populations represented as fraction of total cells. Statistical analysis performed using unpaired t-tests. F) CELESTA plots of CD8+ T cells and EGFR+ macrophages. G) Zoomed in representation of increased anti-colocation of CD8+ T cells and EGFR+ macrophages in tumors. H) Normalized CLQs between CD8+ T cells and EGFR+ macrophages.

**Supplemental Figure 12.**
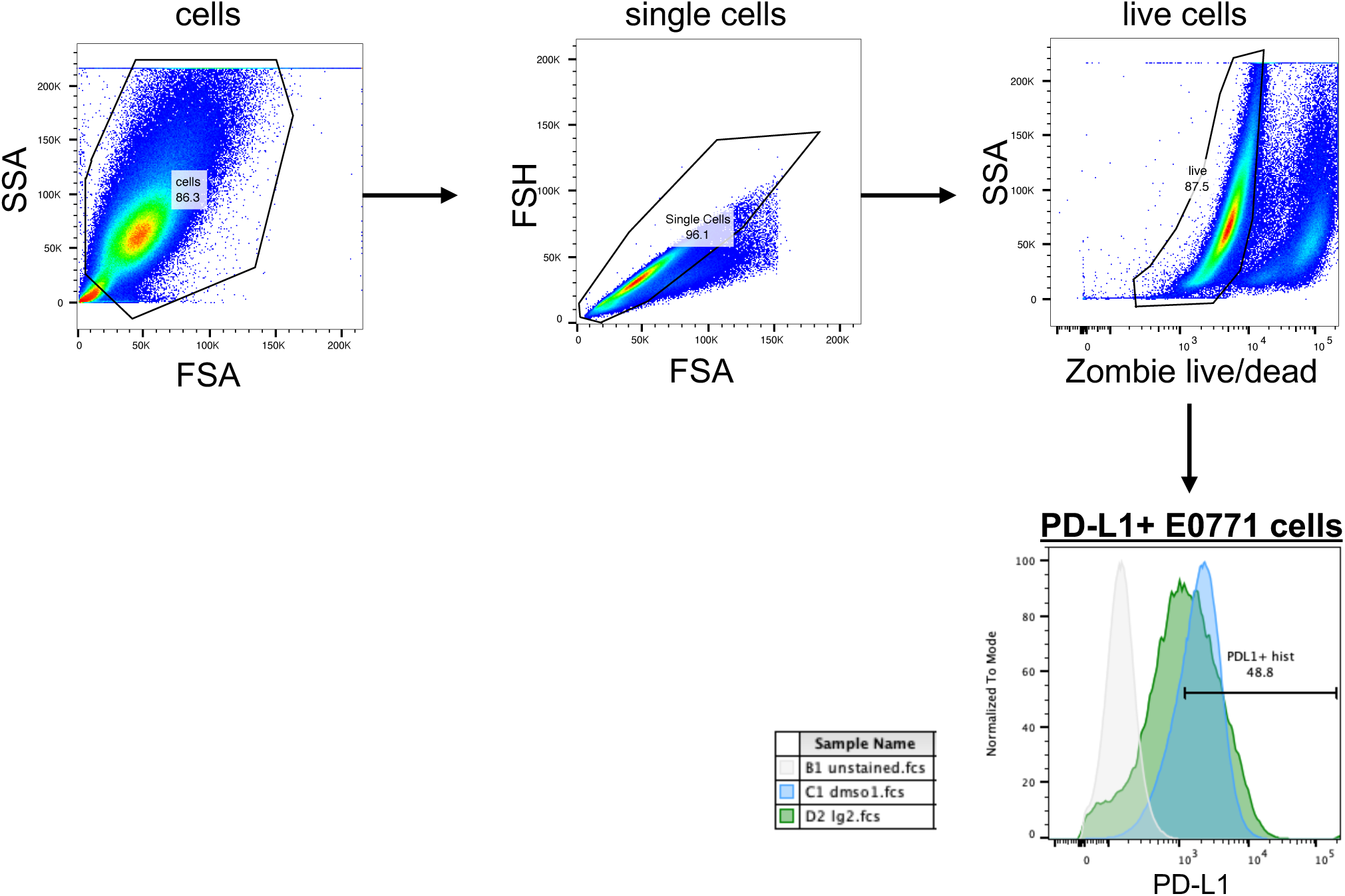
Gating strategy used to examine PD-L1 expression on E0771 cells treated with DMSO or LG1-34. Gating strategy used to identify populations in Fig. 5N-P.

**Supplemental Figure 13.**
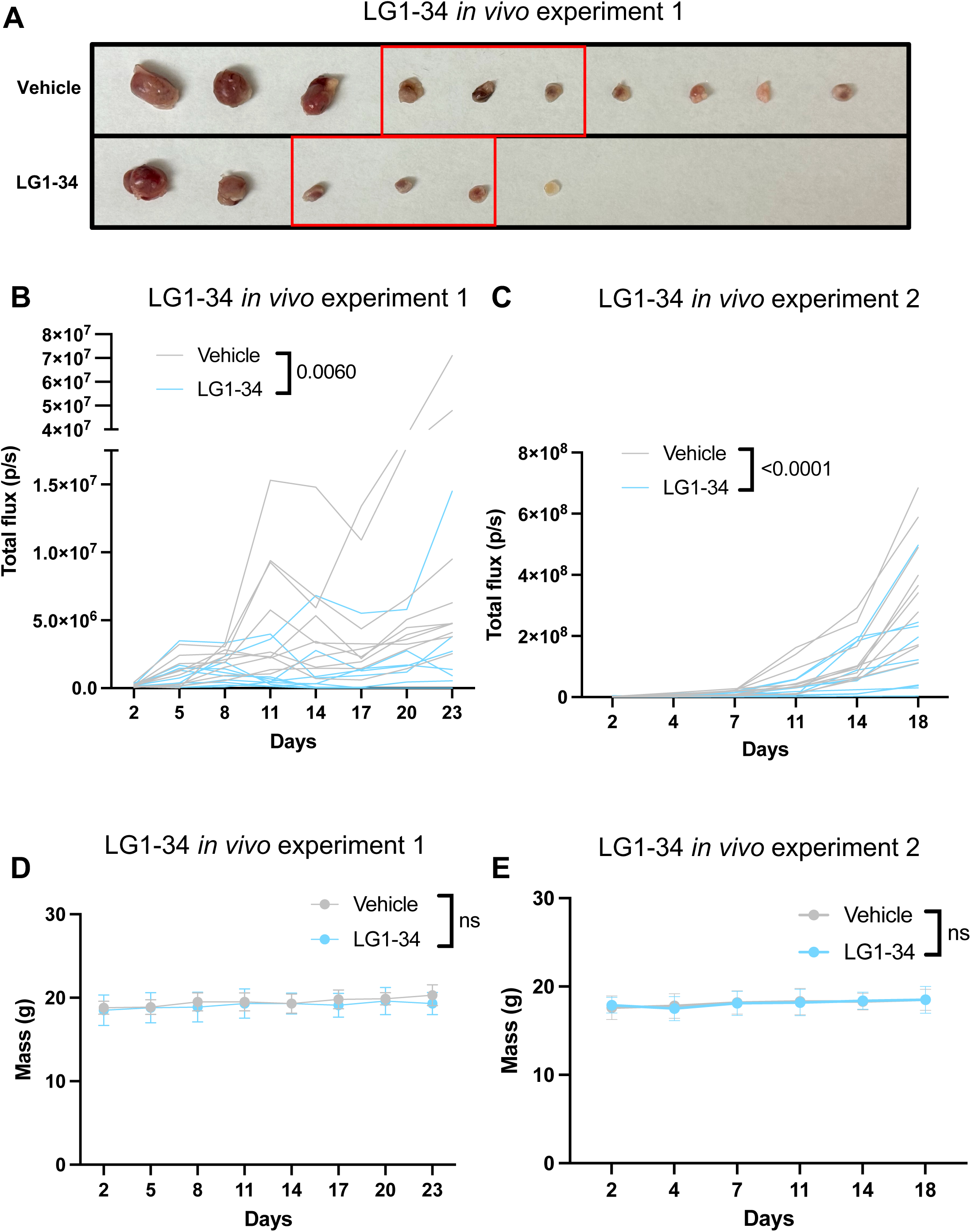
Additional plots for vehicle vs LG1-34 TNBC tumor experiments. A) LG1-34 TNBC experiment #1: Tumors resected on day 23; complete response was observed in 4/10 tumors in LG1-34 treated group; thus only the 6 left-most tumors per group were analyzed via IHC-based spatial analysis. Red boxes surround “size-matched small” tumors analyzed in Fig. 6G, J-K. B) Tumor growth tracks as measured using calipers for individual mice (vehicle n = 10, LG1-34 n = 10) from LG1-34 experiment #1. C) Tumor growth tracks as measured using calipers for individual mice (vehicle n = 10, LG1-34 n = 10) from LG1-34 experiment #2. Mouse weights over time for LG1-34 experiment #1 (D) and #2 (E). Statistical analysis was performed using 2way-ANOVA.

**Supplemental Figure 14.**
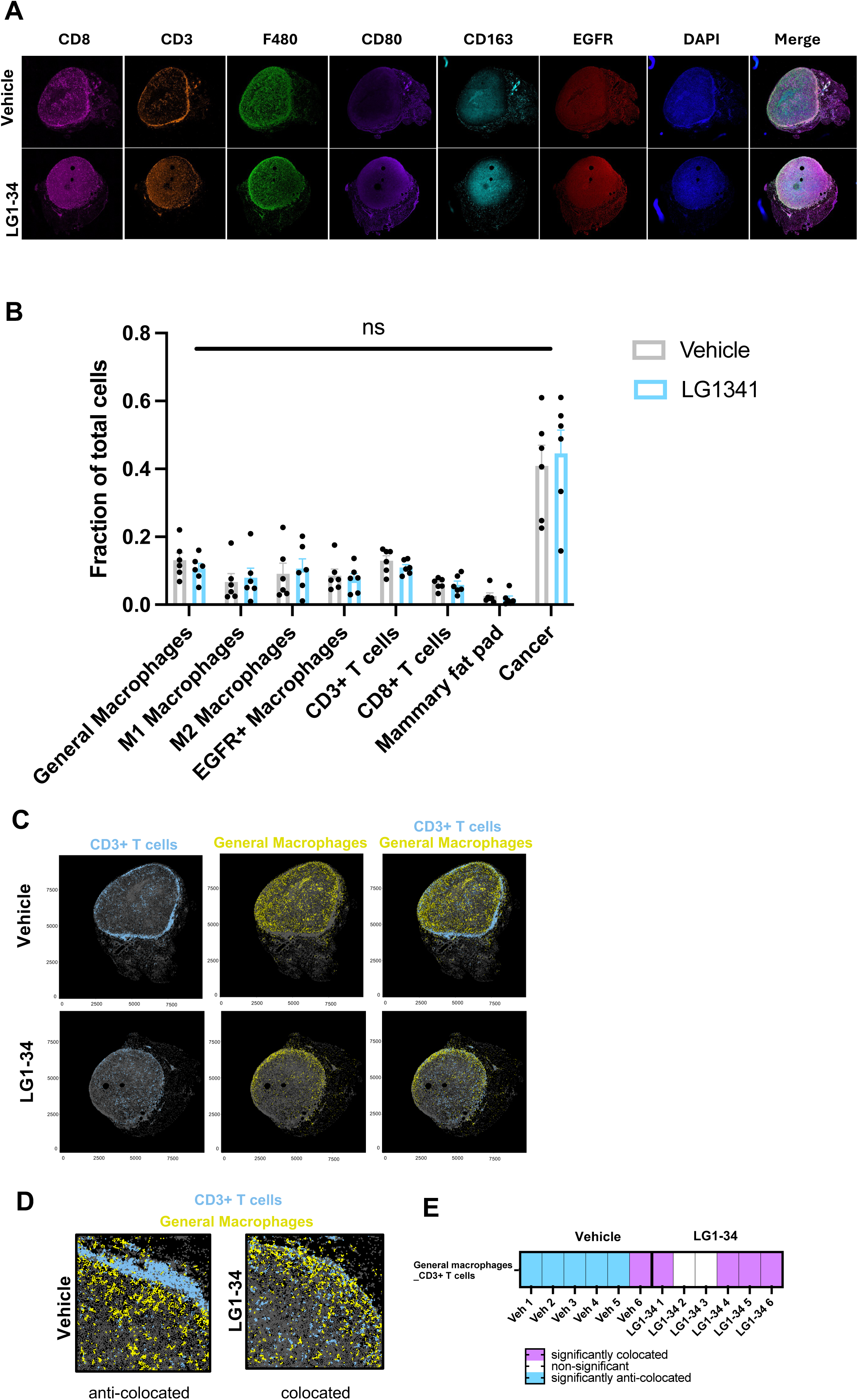
Supplementary material for spatial analysis of LG1-34 treated tumors. A) Fluorescent IHC images of all markers used to identify cell populations through CELESTA. B) Cell populations represented as fraction of total cells within the tumor. Statistical analysis performed using t-tests. C) CELESTA plots of CD3+ T cells and general macrophages. D) Zoomed in representation of colocation between CD3+ T cells (CD3+/CD8-) and general macrophages (F480+ only) in tumors from vehicle and LG1-34 treated mice. E) Colocation analysis of general macrophages and CD3+ T cells in tumors from vehicle and LG1-34 treated mice. Purple = significant colocation within a sample, white = non-significant, blue = significant anti-colocation within a sample. Significance assessed through spatial permutation analysis[60].

**Supplemental Figure 15.**
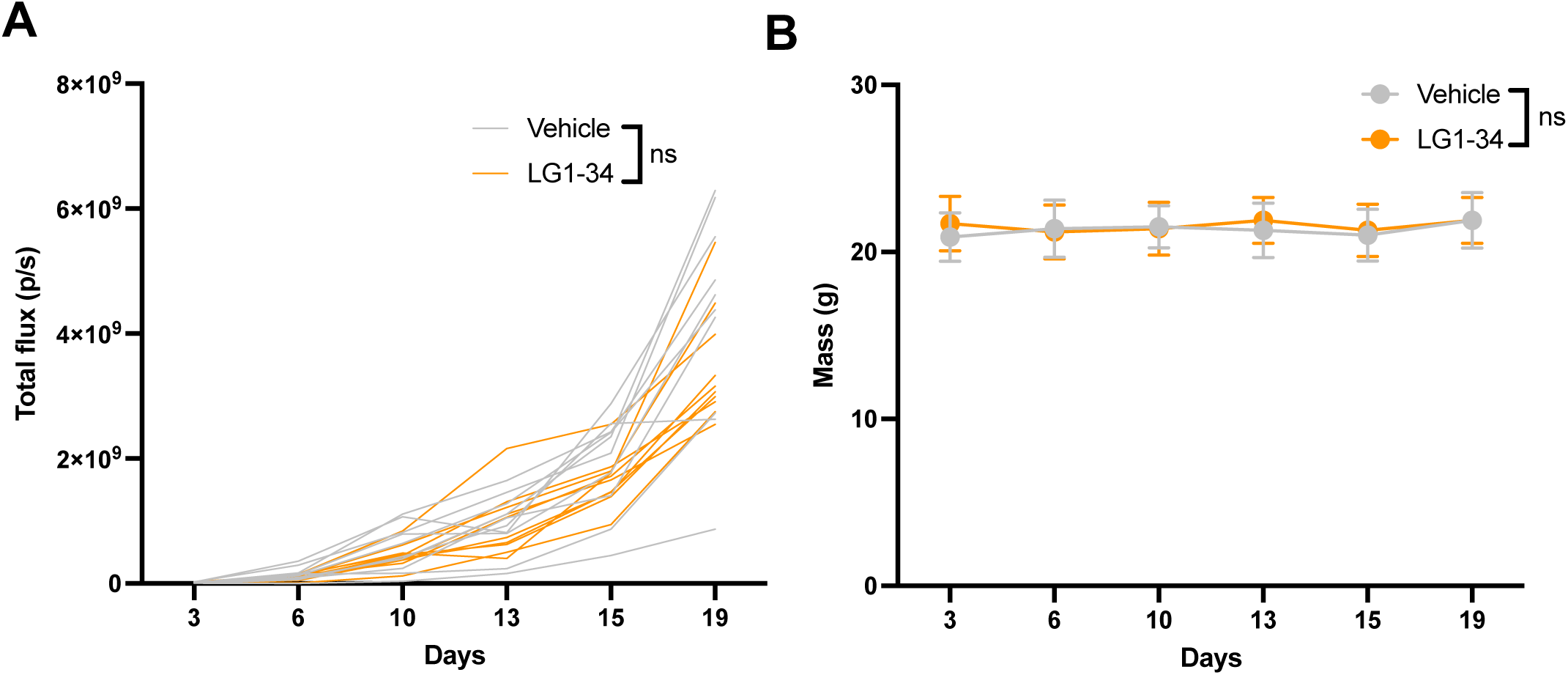
Supplementary plots for Vehicle vs. LG1-34 TNBC experiment in immune-compromised animals. *A*) Tumor growth tracks as measured using IVIS imaging for individual mice from Fig. 6M (vehicle n = 10, LG1-34 n = 10). E) Mouse weights over time. Statistical analysis was performed using 2way-ANOVA.

**Supplemental Figure 16.**
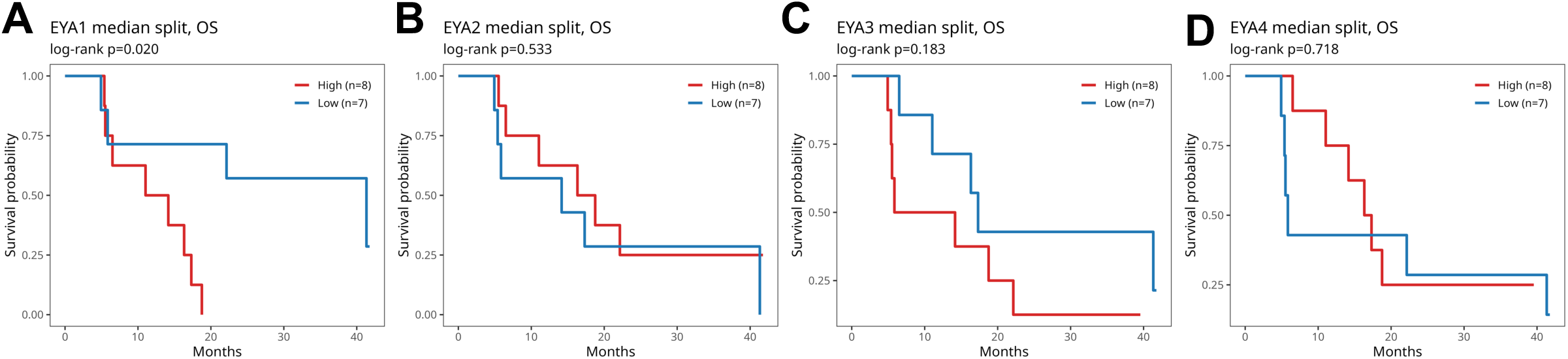
Individual Kaplan-Meier survival plots for overall TNBC survival following pembrolizumab + carboplatin/nab-paclitaxel treatment stratified by Eya1-4 expression level (high/low median split). Kaplan-Meier plots generated from A) Eya1, B) Eya2, C) Eya3, and D) Eya4 expression. Data analyzed from GSE241876.

**Supplemental Figure 17.**
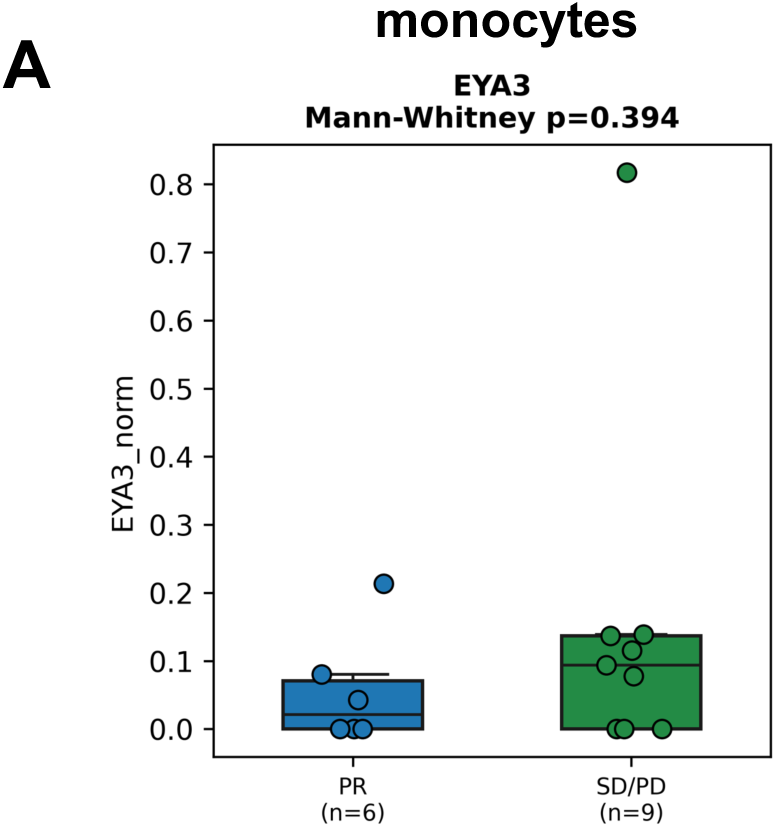
Eya3 mRNA expression within monocytes in TNBC patients with PR or SD/PD following atezolizumab (anti-PD-L1) plus paclitaxel treatment. A) Eya3 expression in monocytes. Data analyzed from GSE169246.

**Supplementary Table 1.** Matrix of IHC markers used to identify different cell populations. Matrix used in combination with CELESTA for identification of cell populations within the tumor immune microenvironment.

| Subpopulation | Lineage level | DAPI | CD163 | CD3 | CD80 | CD8 | EGFR | F480 |
| --- | --- | --- | --- | --- | --- | --- | --- | --- |
| Mammary fat pad | 1_0_1 | 1 | 0 | 0 | 0 | 0 | NA | 0 |
| General Macrophages | 1_0_2 | 1 | 0 | 0 | NA | 0 | NA | 1 |
| M1 Macrophages | 1_0_3 | 1 | 0 | 0 | 1 | 0 | NA | 1 |
| M2 Macrophages | 1_0_4 | 1 | 1 | 0 | 1 | 0 | NA | 1 |
| CD3+ T cells | 1_0_5 | 1 | 0 | 1 | 0 | 0 | 0 | NA |
| CD8+ T cells | 1_0_6 | 1 | 0 | 1 | 0 | 1 | 0 | NA |
| Cancer | 1_0_7 | 1 | 0 | 0 | NA | 0 | NA | NA |
| Hybrid Macrophage | 1_0_8 | 1 | NA | 0 | NA | 0 | 1 | 1 |

**Supplementary Table 2.** Expression Thresholds for EYA3 KO CELESTA. Matrix of thresholds for analysis of tumors from Eya3fl/fl and Eya3fl/fl;MafB-Cre mice (Fig. 4, Supp. Fig 13).

|  |  | Mammary fat<br>pad | General<br>Macrophages | M1<br>Macrophages | M2<br>Macrophages | CD3+ T<br>cells | CD8+ T<br>cells | Cancer | Hybrid<br>Macrophage |
| --- | --- | --- | --- | --- | --- | --- | --- | --- | --- |
| Control_3_1 | high_expression_threshold_anchor | 0.2 | 0.6 | 0.6 | 0.7 | 0.6 | 0.7 | 0.6 | 0.6 |
|  | high_expression_threshold_index | 0.1 | 0.2 | 0.2 | 0.2 | 0.2 | 0.2 | 0.2 | 0.2 |
| Control_3_2 | high_expression_threshold_anchor | 0.2 | 0.6 | 0.6 | 0.7 | 0.6 | 0.7 | 0.6 | 0.6 |
|  | high_expression_threshold_index | 0.1 | 0.2 | 0.2 | 0.2 | 0.2 | 0.2 | 0.2 | 0.2 |
| Control_4_1 | high_expression_threshold_anchor | 0.2 | 0.6 | 0.6 | 0.7 | 0.6 | 0.7 | 0.6 | 0.6 |
|  | high_expression_threshold_index | 0.1 | 0.2 | 0.2 | 0.2 | 0.2 | 0.2 | 0.2 | 0.2 |
| Control_4_2 | high_expression_threshold_anchor | 0.2 | 0.6 | 0.6 | 0.7 | 0.6 | 0.7 | 0.6 | 0.6 |
|  | high_expression_threshold_index | 0.1 | 0.2 | 0.2 | 0.2 | 0.2 | 0.2 | 0.2 | 0.2 |
| Control_5_1 | high_expression_threshold_anchor | 0.2 | 0.6 | 0.6 | 0.7 | 0.6 | 0.7 | 0.6 | 0.6 |
|  | high_expression_threshold_index | 0.1 | 0.2 | 0.2 | 0.2 | 0.2 | 0.2 | 0.2 | 0.2 |
| Control_5_2 | high_expression_threshold_anchor | 0.2 | 0.6 | 0.6 | 0.7 | 0.6 | 0.7 | 0.6 | 0.6 |
|  | high_expression_threshold_index | 0.1 | 0.2 | 0.2 | 0.2 | 0.2 | 0.2 | 0.2 | 0.2 |
| Control_6_1 | high_expression_threshold_anchor | 0.2 | 0.6 | 0.6 | 0.7 | 0.6 | 0.7 | 0.7 | 0.6 |
|  | high_expression_threshold_index | 0.1 | 0.2 | 0.2 | 0.2 | 0.2 | 0.2 | 0.2 | 0.2 |
| Control_6_2 | high_expression_threshold_anchor | 0.2 | 0.6 | 0.6 | 0.7 | 0.6 | 0.7 | 0.7 | 0.6 |
|  | high_expression_threshold_index | 0.1 | 0.2 | 0.2 | 0.2 | 0.2 | 0.2 | 0.2 | 0.2 |
| Eya3_cKO_2_1 | high_expression_threshold_anchor | 0.2 | 0.6 | 0.6 | 0.6 | 0.6 | 0.5 | 0.6 | 0.6 |
|  | high_expression_threshold_index | 0.1 | 0.2 | 0.2 | 0.2 | 0.2 | 0.2 | 0.2 | 0.2 |
| Eya3_cKO_2_2 | high_expression_threshold_anchor | 0.2 | 0.6 | 0.6 | 0.6 | 0.6 | 0.5 | 0.6 | 0.6 |
|  | high_expression_threshold_index | 0.1 | 0.2 | 0.2 | 0.2 | 0.2 | 0.2 | 0.2 | 0.2 |
| Eya3_cKO_7_1 | high_expression_threshold_anchor | 0.2 | 0.6 | 0.6 | 0.7 | 0.6 | 0.5 | 0.6 | 0.6 |
|  | high_expression_threshold_index | 0.1 | 0.2 | 0.2 | 0.2 | 0.2 | 0.2 | 0.2 | 0.2 |
| Eya3_cKO_7_2 | high_expression_threshold_anchor | 0.2 | 0.6 | 0.6 | 0.6 | 0.6 | 0.5 | 0.6 | 0.6 |
|  | high_expression_threshold_index | 0.1 | 0.2 | 0.2 | 0.2 | 0.2 | 0.2 | 0.2 | 0.2 |
| Eya3_cKO_8_1 | high_expression_threshold_anchor | 0.2 | 0.6 | 0.6 | 0.6 | 0.6 | 0.5 | 0.7 | 0.6 |
|  | high_expression_threshold_index | 0.1 | 0.2 | 0.2 | 0.2 | 0.2 | 0.2 | 0.3 | 0.2 |
| Eya3_cKO_8_2 | high_expression_threshold_anchor | 0.2 | 0.6 | 0.6 | 0.6 | 0.6 | 0.5 | 0.7 | 0.6 |
|  | high_expression_threshold_index | 0.1 | 0.2 | 0.2 | 0.2 | 0.2 | 0.2 | 0.3 | 0.2 |
| Eya3_cKO_9_1 | high_expression_threshold_anchor | 0.2 | 0.6 | 0.6 | 0.6 | 0.5 | 0.5 | 0.6 | 0.6 |
|  | high_expression_threshold_index | 0.1 | 0.2 | 0.2 | 0.2 | 0.2 | 0.2 | 0.2 | 0.2 |
| Eya3_cKO_9_2 | high_expression_threshold_anchor | 0.2 | 0.6 | 0.6 | 0.7 | 0.6 | 0.5 | 0.6 | 0.6 |
|  | high_expression_threshold_index | 0.1 | 0.2 | 0.2 | 0.2 | 0.2 | 0.2 | 0.2 | 0.2 |

**Supplementary Table 3.** Expression Thresholds for LG1-34 CELESTA. Matrix of thresholds for analysis of tumors from C57BL/6 mice treated with vehicle or LG1-34 (Fig. 6, Supp. Fig 16).

|  |  | Mammary fat pad | General Macrophages | M1 Macrophages | M2 Macrophages | CD3+ T cells | CD8+ T cells | Cancer | Hybrid Macrophage |
| --- | --- | --- | --- | --- | --- | --- | --- | --- | --- |
| Vehicle_1 | high_expression_threshold_anchor | 0.7 | 0.8 | 0.4 | 0.4 | 0.5 | 0.4 | 0.3 | 0.6 |
|  | high_expression_threshold_index | 0.7 | 0.3 | 0.2 | 0.2 | 0.2 | 0.2 | 0 | 0.2 |
| Vehicle_2 | high_expression_threshold_anchor | 0.7 | 0.8 | 0.4 | 0.4 | 0.5 | 0.4 | 0.3 | 0.6 |
|  | high_expression_threshold_index | 0.7 | 0.3 | 0.2 | 0.2 | 0.2 | 0.2 | 0 | 0.2 |
| Vehicle_3 | high_expression_threshold_anchor | 0.7 | 0.8 | 0.4 | 0.4 | 0.4 | 0.7 | 0.3 | 0.6 |
|  | high_expression_threshold_index | 0.7 | 0.3 | 0.2 | 0.2 | 0.2 | 0.2 | 0 | 0.2 |
| Vehicle_4 | high_expression_threshold_anchor | 0.7 | 0.8 | 0.4 | 0.4 | 0.5 | 0.4 | 0.3 | 0.6 |
|  | high_expression_threshold_index | 0.7 | 0.3 | 0.2 | 0.2 | 0.2 | 0.2 | 0 | 0.2 |
| Vehicle_5 | high_expression_threshold_anchor | 0.7 | 0.8 | 0.4 | 0.4 | 0.4 | 0.7 | 0.3 | 0.6 |
|  | high_expression_threshold_index | 0.7 | 0.3 | 0.2 | 0.2 | 0.2 | 0.2 | 0 | 0.2 |
| Vehicle_6 | high_expression_threshold_anchor | 0.7 | 0.8 | 0.4 | 0.4 | 0.5 | 0.4 | 0.3 | 0.6 |
|  | high_expression_threshold_index | 0.7 | 0.3 | 0.2 | 0.2 | 0.2 | 0.2 | 0 | 0.2 |
| LG1-34_1 | high_expression_threshold_anchor | 0.7 | 0.8 | 0.4 | 0.4 | 0.5 | 0.4 | 0.3 | 0.6 |
|  | high_expression_threshold_index | 0.7 | 0.3 | 0.2 | 0.2 | 0.2 | 0.2 | 0 | 0.2 |
| LG1-34_2 | high_expression_threshold_anchor | 0.7 | 0.8 | 0.4 | 0.4 | 0.4 | 0.7 | 0.3 | 0.6 |
|  | high_expression_threshold_index | 0.7 | 0.3 | 0.2 | 0.2 | 0.2 | 0.2 | 0 | 0.2 |
| LG1-34_3 | high_expression_threshold_anchor | 0.7 | 0.8 | 0.4 | 0.4 | 0.4 | 0.7 | 0.3 | 0.6 |
|  | high_expression_threshold_index | 0.7 | 0.3 | 0.2 | 0.2 | 0.2 | 0.2 | 0 | 0.2 |
| LG1-34_4 | high_expression_threshold_anchor | 0.7 | 0.8 | 0.4 | 0.4 | 0.5 | 0.4 | 0.3 | 0.6 |
|  | high_expression_threshold_index | 0.7 | 0.3 | 0.2 | 0.2 | 0.2 | 0.2 | 0 | 0.2 |
| LG1-34_5 | high_expression_threshold_anchor | 0.7 | 0.8 | 0.4 | 0.4 | 0.5 | 0.4 | 0.3 | 0.6 |
|  | high_expression_threshold_index | 0.7 | 0.3 | 0.2 | 0.2 | 0.2 | 0.2 | 0 | 0.2 |
| LG1-34_6 | high_expression_threshold_anchor | 0.7 | 0.8 | 0.4 | 0.4 | 0.4 | 0.7 | 0.3 | 0.6 |
|  | high_expression_threshold_index | 0.7 | 0.3 | 0.2 | 0.2 | 0.2 | 0.2 | 0 | 0.2 |

**Supplementary Table 4.** Primary and secondary antibody reagent information. All primary and secondary antibodies and pertinent information used in Western blot assays.

| Primary antibody | Source | Identifier | Clone Number (if Applicable) |
| --- | --- | --- | --- |
| Rabbit polyclonal anti-EYA1 | Proteintech | 22658-1-AP | N/A |
| Rabbit monoclonal anti-EYA2 | Invitrogen | MA5-52842 | 23GB5350 |
| Rabbit polyclonal anti-EYA3 | Bethyl | A302-689A | N/A |
| Mouse monoclonal anti-EYA4 | Santa Cruz Biotech | sc-393111 | E-11 |
| Rabbit monoclonal anti-GAPDH | Cell Signaling Technology | 2118S | 14C10 |
| Mouse monoclonal anti-β-Actin | Sigma-Aldrich | A5316 | AC-74 |
| Goat anti-mouse HRP Secondary antibody | LICORbio | 926-80010 | N/A |
| Goat anti-rabbit HRP secondary antibody | LICORbio | 92680011 | N/A |

**Supplementary Table 5.** Flow cytometry antibodies. Antibody panels used for flow cytometry analysis of tissues or cells.

T and B cell panel
|  | Marker | Color | Catalog no. | Clone |
| --- | --- | --- | --- | --- |
| 1 | Zombie Live Dead | Aqua | 423101 |  |
| 2 | CD45 | BV421 | 103134 | 30-F11 |
| 3 | CD3 | BV605 | 100237 | 17A2 |
| 4 | CD8a | BV650 | 100742 | 53-6.7 |
| 5 | CD4 | PE | ebio |  |
| 6 | CD19 | APC-Cy7 | 115529 | 6D5 |
| 7 | B220 | PerCPCy5.5 | biol | RA3-6B2 |

NK cell panel
|  | Marker | Color | Catalog no. | Clone |
| --- | --- | --- | --- | --- |
| 1 | Zombie Live Dead | Aqua | 423101 |  |
| 2 | CD45 | BV421 | 103134 | 30-F11 |
| 3 | CD3 | BV605 | 100237 | 17A2 |
| 4 | NKp46 | PE | 137604 | 29A1.4 |
| 5 | CD49b | PE-Cy7 | 108905 | DX5 |
| 6 | CD11b | PerCP Cy5.5 | #45-0112-82 | M1/70 |
| 7 | CD27 | APC-Cy7 | 124225 | LG.3A10 |

Innate Immune Panel (some markers switched in and out to maintain panel compatibility)
|  | Marker | Color | Catalog no. | Clone |
| --- | --- | --- | --- | --- |
| 1 | Zombie Live Dead | Aqua (BV510) | 423101 |  |
| 2 | CD45 | BV421 | 103134 | 30-F11 |
| 3 | CD3 | BV605 | 100237 | 17A2 |
| 4 | F4/80 | FITC | 123108 | BM8 |
| 5 | Ly6C | BV711 | 128037 | HK1.4 |
| 6 | Ly6G | AF700 | 127621 | 1A8 |
| 7 | CD11b | PerCP Cy5.5 | ebio | M1/70 |
| 8 | CD11c | BV785 | 117335 | N418 |
| 9 | MHC-II (I-A/I-E) | APC/ Cy7 | 107628 | M5/114.15.2 |
| 10 | CD206 | PE | 141705 | C068C2 |
| 11 | CD80 | APC | #17-0801-82 | 16-10A1 |
| 12 | PD-L1 | PE-Cy7 | biol | 10F.9G2 |
| 13 | MERTK | APC | Biol |  |
| 14 | iNOS | BUV737 | 367-5920-80 | CXNFT |
| 15 | CD64 | BV605 | 139323 | <u>X54-5/7.1</u> |
| 16 | CD80 alt. | BV650 | 100742 | 53-6.7 |

